# Engineering an Exo70 integrated domain of a barley NLR for improved blast resistance

**DOI:** 10.1101/2024.12.24.630226

**Authors:** Indira Saado, Helen J. Brabham, Josh W. Bennett, Anson Ho Ching Lam, Inmaculada Hernández-Pinzón, Matthew J. Moscou, Juan Carlos De la Concepcion, Mark J. Banfield

**Author notes:** Corresponding authors: Juan Carlos De la Concepcion, Mark J. Banfield.

## Abstract

Intracellular immune receptors protect plants from microbial invasion by detecting and responding to pathogen-derived effector molecules, often triggering cell death responses. However, pathogen effectors can evolve to avoid immune recognition, resulting in devastating diseases that threaten global agriculture. Here, we show that an integrated Exo70 domain from the barley NLR RGH2 can interact with both the rice blast pathogen effector AVR-Pii and a closely related wheat blast variant. We used structure-led engineering to develop RGH2^+^ that shows increased binding affinity towards AVR-Pii variants and increased cell death responses on heterologous expression in *Nicotiana benthamiana*. Infection assays in transgenic barley lines carrying RGH2^+^ with the paired NLR RGH3 indicate a reduced susceptibility to blast strains expressing AVR-Pii variants. These results demonstrate the potential of engineering NLR receptors as an effective strategy for improving resistance towards one of the most destructive diseases affecting cereal production.

## Introduction

Plants are continuously threatened by pathogens and rely on an innate immune system for defence. One critical component of this immune system is the recognition of virulence proteins (effectors) that are delivered from pathogens to manipulate host cells for the benefit of the invading organism. *Magnaporthe oryzae* is a devastating ascomycete fungus that infects cereal crops, including rice, wheat, barley, and other grasses, causing blast disease^1,2^. Since the pathogen’s host jump to wheat in South America in the 1980s^3,4^, and recent spread to Asia and Africa^5^, wheat blast disease (caused by *M. oryzae* pathotype *Triticum*) has emerged as major pandemic threat to agriculture. *M. oryzae* deploys a diverse set of effectors during colonization, targeting key plant processes to promote infection. Despite their essential roles in infection, the function of only a limited number of *M. oryzae* effectors has been fully characterized.

Pathogen effectors can be sensed by intracellular nucleotide-binding leucine-rich repeat (NLR) receptors, triggering immune responses that often result in localised cell death, limiting the spread of infection. As such, engineering NLRs for novel recognition profiles can unlock promising strategies for improved disease resistance^6–9^. Canonical NLRs comprise an N-terminal signalling domain, a central nucleotide-binding adaptor NB-ARC switch domain, and a C-terminal leucine-rich repeat (LRR) domain involved in intramolecular molecular contacts and pathogen effector recognition^10^. While some NLRs function alone to both sense and respond to effectors (singleton NLRs), others function in pairs, with a sensor NLR to recognize the effector, directly or indirectly, and a helper NLR required for immune response execution. Such pairs can be co-located in plant genomes including in head-to-head orientations with shared promoters, or function as part of distributed genetic networks^11,12^.

To date, two co-located paired NLR systems have been extensively studied in rice: Pik-1/Pik-2^13,14^ and RGA5/4^15,16^. Both sensor NLRs, Pik-1 and RGA5, contain non-canonical integrated heavy metal-associated (HMA) domains to bait specific effectors from *M. oryzae*^17–20^. The sensor NLR Pik-1 contains a HMA domain positioned between the CC and NB-ARC domains, and with the helper NLR Pik-2 cooperatively triggers immunity against the blast pathogen effector AVR-Pik^13,14,18,21,22^. The Pik/AVR-Pik system has emerged as a model for how receptor engineering can lead to novel immune outcomes^6–8,19,20,23,24^. By contrast, the sensor NLR RGA5 has an HMA domain at the C-terminus (after the LRR) and baits the blast pathogen effectors AVR-Pia and AVR1-CO39^25^. Activation of immunity requires the helper NLR RGA4^26,27^. This NLR pair has also been leveraged for engineering immunity to expand effector recognition profiles^28^.

In addition to direct effector recognition, paired NLRs can indirectly recognise pathogen effectors^29–31^. For example, the unconventional RIN4 (RPM1-Interacting Protein 4)-like NOI (nitrate induced) domain of the rice sensor NLR Pii-2 monitors the status of host OsExo70-F2/3 proteins targeted by the rice blast Zinc-finger fold (ZiF) effector AVR-Pii^29–32^. Canonical Exo70 proteins are part of the exocyst, a conserved protein complex essential for tethering secretory vesicles to membranes^33,34^.However, in plants, the Exo70 family has dramatically expanded^35^ and acquired novel functions independent from the exocyst^33,36–38^, including plant defence^39–43^, making this protein a target for pathogen effectors^44^. Integrated domains in NLRs, such as HMAs, are likely derived from the host virulence-associated targets of effectors, repurposed for effector perception^45^. Recently, a head-to-head NLR pair from the barley cultivar Baronesse was identified comprising the sensor NLR RGH2, containing a C-terminal Exo70 integrated domain, and its helper NLR RGH3^46^. The integration of an Exo70 domain in RGH2^47^ suggests a potential role in immunity by serving as an integrated domain to bait effectors for recognition.

In this study, we show that the Exo70 integrated domain of the barley sensor NLR RGH2 can interact with the rice blast effector AVR-Pii and a closely related wheat blast variant in vitro and in vivo. On co-expression with the helper NLR RGH3, the RGH2/RGH3 pair elicits limited cell death to the AVR-Pii variants in *N. benthamiana*. Based on the crystal structure of the OsExo70F2/AVR-Pii complex^29^, we engineered a new immune receptor, RGH2^+^, that showed enhanced binding affinity towards AVR-Pii variants, leading to improved cell death responses on co-expression with RGH3 in *N. benthamiana*. In transgenic barley, RGH2^+^/RGH3 confers a reduction in disease severity compared to the wild-type RGH2/RGH3 when infected with *M. oryzae* expressing AVR-Pii. These findings establish the RGH2/RGH3 NLR pair as a platform for engineering enhanced effector response profiles and highlights new opportunities for protecting cereal crops from blast disease.

## Results

### Multiple Exo70 domains are integrated in NLRs across the Poales order

Previous work established that NLRs from diverse plant species carry integrated domains^48,49^. We sought to identify NLRs with integrated Exo70 domains to understand their source and assess their use as scaffolds for NLR engineering. To enrich the set of NLRs for analysis, we performed de novo transcriptome assembly of publicly available RNAseq data for 102 Poales species including a total of 292 accessions (Fig. 1A). Domain annotation identified 184,775 NB-ARC domain containing proteins. Within this set of NB-ARC containing proteins, 82 were found to also have an Exo70 domain. Comparative analysis of these integrated Exo70 domains found that the 82 putative integrated Exo70 domains are derived from eight unique fusion events. Among these, six domains were found to be of sufficient length for phylogenetic analysis with grass Exo70 domains^43^. Maximum likelihood phylogenetic analysis using Exo70 from eight grass species found the six integrated Exo70 domains were placed in either the Exo70F or Exo70FX clades (Fig. 1B). The most prevalent integration is RGH2 that carries a C-terminal fusion with Exo70F1, which was originally identified at the *Mla* locus in the barley (*Hordeum vulgare*) accession Baronesse. This integration event was found in seven species within Triticeae and Poeae tribes^47^. In barley, a second NLR, Hvu23934, was found with integration of Exo70FX2 in three Triticeae species (*Aegilops sharonensis*, *Ae. speltoides*, and *H. vulgare*). The remaining four integrated domains are found in single species: *Arundinella pubescens* (Panicoideae; Apu26724; NLR-Exo70F4), *A. pubescens* (Panicoideae; Apu28443; NLR-Exo70F2), *Holcus lanatus* (Pooideae; Hla80948; NLR-Exo70FX2), and *Juncus effusus* (Juncaceae; Jef9120; NLR-Exo70F) (Supplementary Fig. S1). These integrated domains (excluding Apu26724) were taken forward for further analysis.

**Fig. 1:**
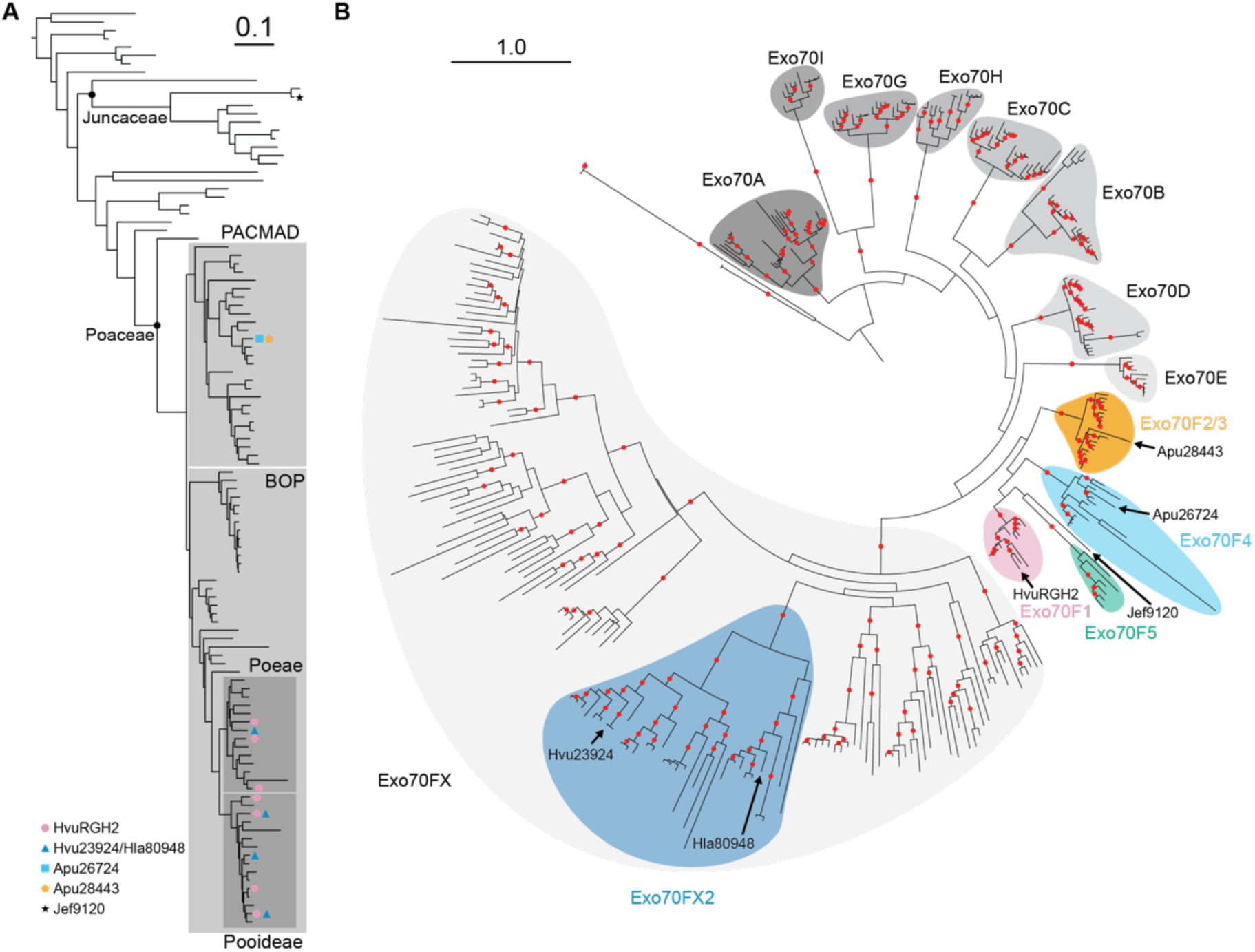
NLRs with integrated Exo70 domains in the Poales are derived from the Exo70F and Exo70FX clades. **A.** Maximum likelihood phylogenetic tree for Poales species based on 1,019 Benchmarking Universal Single-Copy Ortholog (BUSCO) genes from 110 species spanning 1,471,281 variable sites. Outgroup species include oilpalm (*Elaeis guineensis*) and banana (*Musa acuminata*). Legend indicates the presence of integrated Exo70 domains using coloured coded shapes. Important families, clades, and subfamilies in the Poales carrying integrated Exo70 domains are indicated in shaded boxes. Scale indicates substitutions per site. **B.** Maximum likelihood phylogenetic tree of grass Exo70 proteins showing that integrated Exo70 domains are derived from the Exo70F and Exo70FX clades. Grass species include barley (*Hordeum vulgare*), wheat (*Triticum aestivum*), purple false brome (*Brachypodium distachyon*), rice (*Oryza sativa*), *Oropetium thomaeum*, maize (*Zea mays*), foxtail millet (*Setaria italica*), and sorghum (*Sorghum bicolor*). Exemplar integrated Exo70 domains include *H. vulgare* RGH2 (Exo70F1) and Hvu23934 (Exo70FX2), *Arundinella pubescens* Apu26724 (Exo70F4) and Apu28443 (Exo70F2), *Holcus lanatus* Hla80948 (Exo70FX2), and *Juncus effusus* Jef9120 (unclassified Exo70F). Outgroup Exo70s include *Saccharomyces cerevisiae* (yeast) and *Mus musculus* (mouse). Scale indicates substitutions per site.

### Rice blast AVR-Pii and the wheat blast variant bind the barley RGH2-Exo70 integrated domain in yeast

As some ZiF effectors, including AVR-Pii, bind rice Exo70s from clade F, we tested whether Exo70 integrated domains could bind ZiF effectors from rice and wheat-infecting blast isolates of *M. oryzae* in pairwise yeast two-hybrid (Y2H) assays. For this we cloned the Exo70 integrated domains from the NLRs RGH2, Hvu23924, Hla80948, Jef9120, and Apu28443 and tested their binding to representatives of each ZiF effector clade from rice and wheat-infecting blast isolates. These assays revealed that ZiF effectors from tribe VIII, from both rice blast (ZiF_VIIId, hereafter AVR-Pii^Rb^) and wheat blast (ZiF_VIIIc, hereafter AVR-Pii^Wb^) isolates bound RGH2-Exo70, as did tribe IX member ZiF_IXj (Fig. 2A). We did not observe interactions between other NLR Exo70 integrated domains and any ZiF effectors tested (Supplementary Fig. S3A, S4A, S5A). We observed yeast growth with Apu28443-Exo70 in the presence of all tested ZiF effectors, likely indicating autoactivity (Supplementary Fig. S6A). Therefore, we did not investigate it further in this study. Protein accumulation was assessed by western blot (Supplementary Fig. S2A, S3B, S4B, S5B, S6B).

**Fig. 2:**
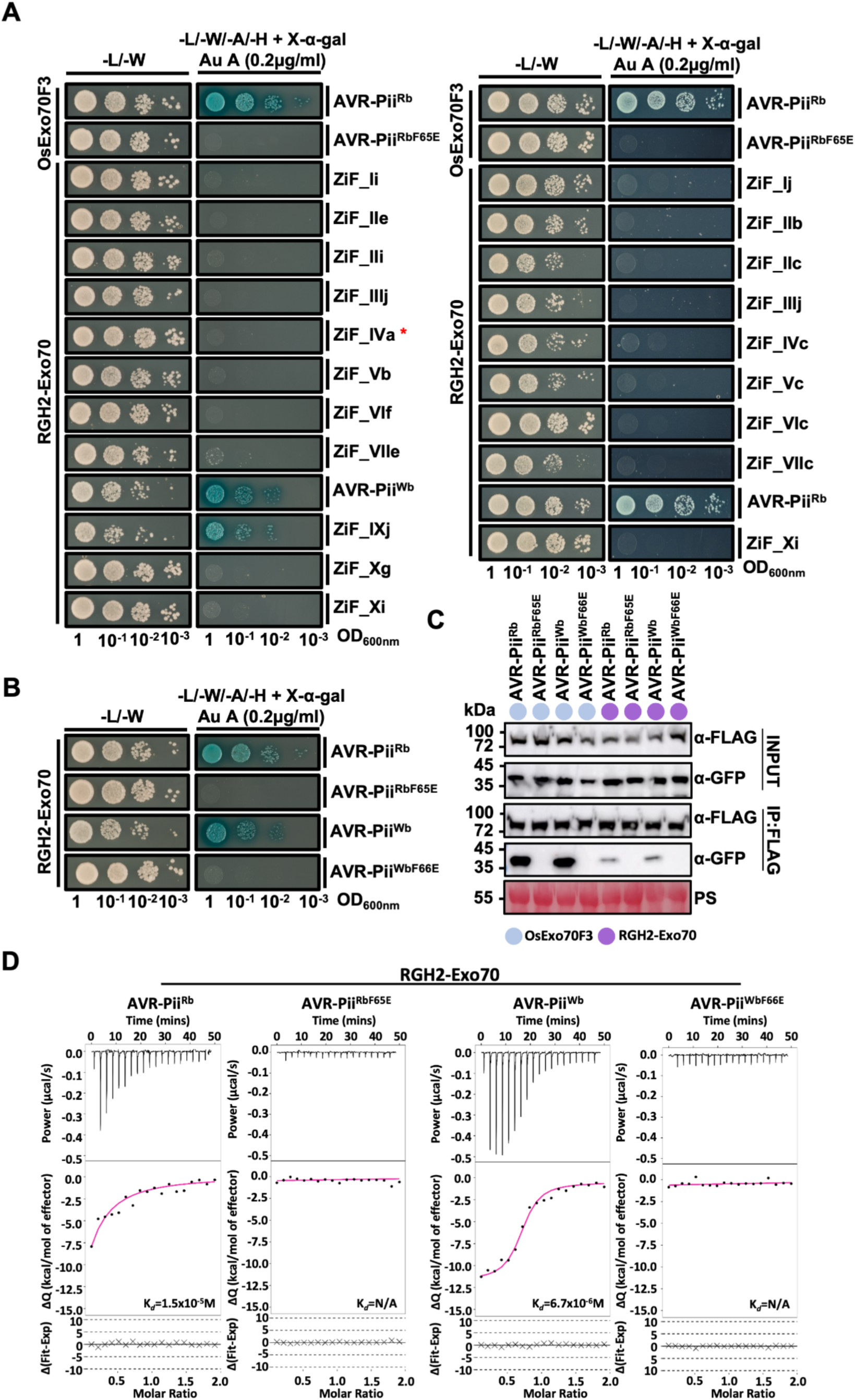
RGH2 interacts with rice blast AVR-Pii and the wheat blast variant in vitro and in vivo. **A.** Y2H binding assay of the most prevalent ZiF effectors from wheat (left) and rice (right) blast isolates as previously defined^32^ to the RGH2-Exo70 integrated domain. Yeast cells transformed with both plasmids were spotted onto selective synthetic dropout (SD) media. For plates with quadruple-dropout media X-α-Gal and Aureobasidin A (Au A) were added to the media. Yeast growth was monitored for four days after plating. Growth and development of blue coloration indicates protein-protein interactions. The experiment was repeated three times with comparable results. **B.** Interaction of rice blast AVR-Pii, the wheat blast variant and mutants with RGH2-Exo70 using the Y2H assay. The experiment was carried out as described for panel A. Yeast growth was monitored for four days subsequent to plating. The experiment was repeated a minimum of three times, with similar results. **C.** Co-IP assay showing that rice blast AVR-Pii and the wheat blast variant interact with RGH2-Exo70-FLAG in *N. benthamiana.* The co-IP assays used an anti-FLAG antibody to pull down AVR-Pii variants. Immunoblot analysis using an anti-GFP antibody shows that RGH2-Exo70-FLAG and OsExo70F3-FLAG co-purified with rice and wheat blast GFP-AVR-Pii variants but not with the effector mutants. Ponceau S staining shows the total amount of protein loaded per lane in the input. The experiment was repeated three times with similar results. **D.** Binding of AVR-Pii variants to RGH2-Exo70 in vitro, as determined by ITC. Upper panel shows the heat differences upon injection of AVR-Pii variants and mutants into the cell containing RGH2-Exo70. Middle panel shows the integrated heats of injection (dots-black colour) and best fit (solid line-magenta colour) to a single-site binding model calculated using AFFINImeter ITC analysis software^68^. Bottom panel indicates the difference between the fit to a single-site binding model and the experimental data. The experiments shown are representative of three replicates.

The crystal structure of AVR-Pii^Rb^ in complex with OsExo70F2^29^ identified Phe-65 as a key interacting residue from the effector, and a Phe65Glu mutation in AVR-Pii^Rb^ abolished the interaction with OsExo70F3^29^. AVR-Pii^Wb^ is predicted to adopt the conserved ZiF structural fold, as observed for AVR-Pii^Rb^ (Supplementary Fig. S7A, B), and binds to OsExo70F3^32^. To test whether AVR-Pii^Rb^ and AVR-Pii^Wb^ interact with the RGH2-Exo70 integrated domain via a similar interface, we tested the binding of both AVR-Pii^Rb^ and AVR-Pii^Wb^ Phe to Glu mutants to RGH2-Exo70 by Y2H. This assay revealed that these mutants do not bind RGH2-Exo70 (Fig. 2B). Protein accumulation was assessed by western blot (Supplementary Fig. S2B).

### AVR-Pii^Rb^ and AVR-Pii^Wb^ bind the barley RGH2-Exo70 integrated domain in planta

To further validate the interaction between AVR-Pii^Rb^ or AVR-Pii^Wb^ with RGH2-Exo70, we used co-immunoprecipitation (co-IP) assays in *N. benthamiana*. RGH2-Exo70-FLAG was co-expressed with GFP-AVR-Pii^Rb^, GFP-AVR-Pii^Wb^ and their respective Phe to Glu mutants. We also co-expressed the effectors with OsExo70F3-FLAG as positive and negative controls, respectively. Following extraction and immunoprecipitation with anti-FLAG beads (to capture Exo70 domains), both effectors, but not their Phe to Glu mutants, were pulled down (Fig. 2C). Interestingly, we observe that the detected levels of AVR-Pii^Rb^ and AVR-Pii^Wb^ in the co-IPs are higher with OsExo70F3 compared to the RGH2-Exo70, suggestive of a stronger association.

### AVR-Pii^Rb^ and AVR-Pii^Wb^ bind more weakly to RGH2-Exo70 than OsExo70F3 in vitro

To further investigate the binding of AVR-Pii^Rb^ and AVR-Pii^Wb^ to RGH2-Exo70 we produced and purified these proteins from *Escherichia coli*. We then performed isothermal titration calorimetry (ITC) assays to assess the binding strength between effectors and the RGH2-Exo70 domain. Previous results show that AVR-Pii^Rb^ binds to OsExo70F3 in the nano-molar range (*KD* of 1.0×10^−10^ M)^29^. RGH2-Exo70 binds AVR-Pii^Rb^ and AVR-Pii^Wb^ in the micro-molar range (*KD*s of 1.5×10^−5^ M and 6.7×10^−6^ M respectively), showing that AVR-Pii^Rb^ binds OsExo70F3 more tightly than RGH2-Exo70 in vitro (Supplementary Fig. S2D), and AVR-Pii^Wb^ binds RGH2-Exo70 in the same range as AVR-Pii^Rb^. No binding was observed with the effector mutants AVR-Pii^RbF65E^ or AVR-Pii^WbF66E^ with OsExo70F3 or RGH2-Exo70 (Fig. 2D)^29^. Taken together, the results from Y2H, co-IP, and ITC show that while AVR-Pii^Rb^ and AVR-Pii^Wb^ bind to RGH2-Exo70, this interaction is not as robust as to OsExo70F3. This suggests that engineering RGH2-Exo70 for improved effector binding could enhance effector perception and immunity-related signalling by this receptor.

### An engineered RGH2^+^ NLR shows higher binding affinity towards AVR-Pii^Rb^ and AVR-Pii^Wb^

We hypothesised that engineering a new binding interface to RGH2-Exo70, mimicking OsExo70F3, could improve interaction strength. Guided by the structure of the OsExo70F2/AVR-Pii complex^29^, we generated a modified version of RGH2, RGH2^+^, by introducing three mutations, Ala1378Val, Ile1379Leu, and Leu1396Met (Fig. 3A). AlphaFold3^50^ predictions of the RGH2-Exo70 and RGH2^+^-Exo70 structures indicate a conserved fold with OsExo70F2 (Fig. 3B).

**Fig. 3:**
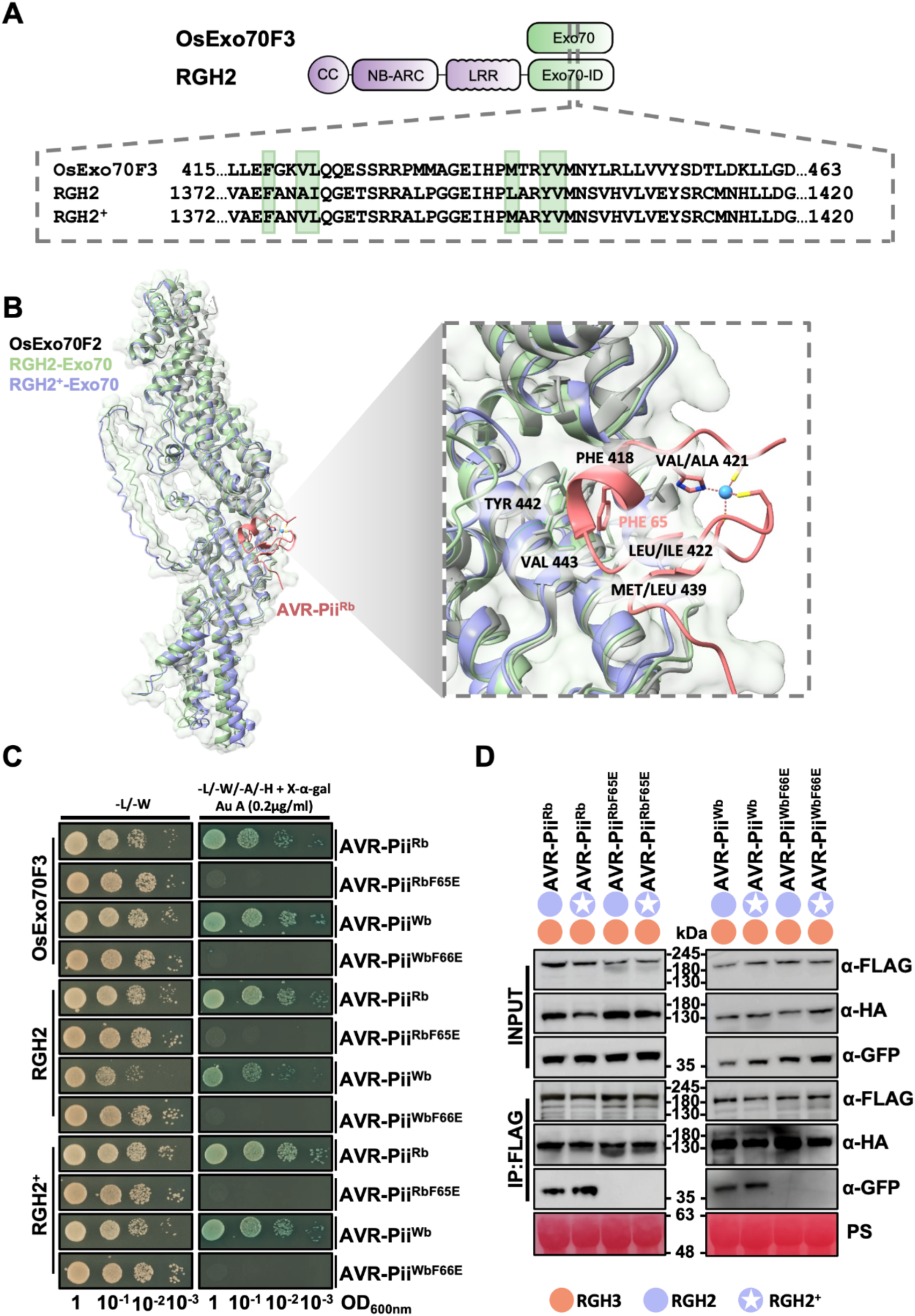
RGH2^+^ binds rice blast AVR-Pii and the wheat blast variant. **A.** Graphical representation of OsExo70F3 and the barley NLR RGH2. Sequence alignment of residues located at the effector binding interface in the OsExo70F2/AVR-Pii complex^29^. Residues important for the formation of the effector binding pocket are highlighted in green. To generate the engineered NLR RGH2^+^, amino acid mutations Ala1378Val, Ile1379Leu, and Leu1396Met were made. **B.** Left, Structural predictions generated with AlphaFold3 as used to interpret interactions between AVR-Pii^Rb^ and RGH2-Exo70/RGH2^+^-Exo70. The structure of OsExo70F2 in complex with AVR-Pii^Rb^ (PDB:7pp2) was previously determined^29^. Right, zoom-in view of the binding interface in the OsExo70F2/AVR-Pii^Rb^ complex, with RGH2-Exo70 and RGH2^+^-Exo70 superimposed. The residues of OsExo70F2 involved in binding the effector that are mutated in this study are indicated (Fig. 3A). The AVR-Pii effector is shown in peach colour, with the Phe65 residue highlighted. **C.** Y2H assay of rice blast AVR-Pii, the wheat blast variant, and mutants with OsExo70F3, barley RGH2-Exo70 and RGH2^+^-Exo70. Left, control plate for yeast growth; right, the quadruple-dropout medium supplemented with X-α-Gal and Aureobasidin A (Au A). The presence of yeast growth and the development of blue coloration in the selection plate signify positive protein-protein interactions. Yeast growth was monitored four days after plating. Each experiment was repeated at least three times, with consistent results obtained. **D.** Co-IP assay showing that rice blast AVR-Pii, and the wheat blast variant, interact with full length RGH2 and RGH2^+^ co-expressed with the helper NLR RGH3 in *N. benthamiana.* Co-IP assays were performed using an anti-FLAG antibody to pull down RGH2-FLAG and RGH2^+^-FLAG interacting proteins. Immunoblot analysis using an anti-GFP antibody showed that GFP-AVR-Pii variants were pulled down with FLAG-RGH2 and FLAG-RGH2^+^, but mutant effectors showed no interaction. Anti-HA was used to confirm the presence of RGH3-HA and interaction with RGH2. RGH2 and RGH2^+^ were only detected weakly in the inputs, but immunoprecipitated proteins are clear. Ponceau S staining shows the amount of total protein loaded per lane in the input. An expanded blot with additional controls is shown in Supplementary Fig.9. Experiment was repeated at least three times, with similar results obtained.

To assess the binding capabilities of the RGH2^+^-Exo70 domain towards AVR-Pii^Rb^ and AVR-Pii^Wb^ we first used the Y2H assay. This showed that the engineered NLR retained binding to these two effectors, and confirmed the same surface of the effector is required for binding as the respective Phe to Glu mutations prevented interaction (Fig. 3C). Protein accumulation was assessed by western blot (Supplementary Fig. S8).

To compare interactions between AVR-Pii^Rb^ and AVR-Pii^Wb^ with RGH2 or RGH2^+^ in plant cells, we first used co-IP. FLAG-tagged full-length RGH2 and RGH2^+^ immunoprecipitated GFP-tagged AVR-Pii^Rb^ and AVR-Pii^Wb^, but not the Phe to Glu mutants of each effector (Fig. 3D). Additional negative controls, including a double mutant Tyr to Ala, Phe to Glu, and a ZiF effector from a tribe that does not interact with Exo70s (ZiF_IIb), were not immunoprecipitated by RGH2 or RGH2^+^ (Supplementary Fig. S9). Secondly, we used the split GAL4 RUBY assay that allows quantification of protein-protein interactions in plant cells through the production of betalain^51^. Agrobacterium strains carrying OsExo70F3-Gal4-BD, RGH2-Exo70-Gal4-BD, or RGH2^+^-Exo70-Gal4-BD were co-infiltrated with constructs containing VP16-tagged AVR-Pii^Rb^ or AVR-Pii^Wb^ or their Phe to Glu mutants (all constructs under the control of the *35Sprom*). Betalain production was observed 3 days post-infiltration (dpi) (Supplementary Fig. S10A, S11A). Intriguingly, significantly higher betalain production was observed in RGH2^+^-Exo70 leaves compared to RGH2-Exo70 with AVR-Pii^Rb^. However, with VP16-AVR-Pii^Wb^, levels of betalain production accumulated at similar levels. Low betalain production was observed with the VP16-tagged effector mutants in combination with Exo70 constructs. To determine whether lower effector expression in *N. benthamiana* leaves could allow distinction between RGH2-Exo70 and RGH2^+^-Exo70 responses to AVR-Pii^Wb^ in this assay, we generated AVR-Pii expression constructs under the control of the *MAS* promoter. Notably, overall betalain production was lower compared to the *35Sprom* assays. However, in these conditions, we observed significant differences between RGH2-Exo70 and RGH2^+^-Exo70 with both AVR-Pii^Rb^ and AVR-Pii^Wb^ (Fig. 4A, B). Protein accumulation for all constructs was assessed by western blot (Supplementary Fig. S10B, S11B).

**Fig. 4:**
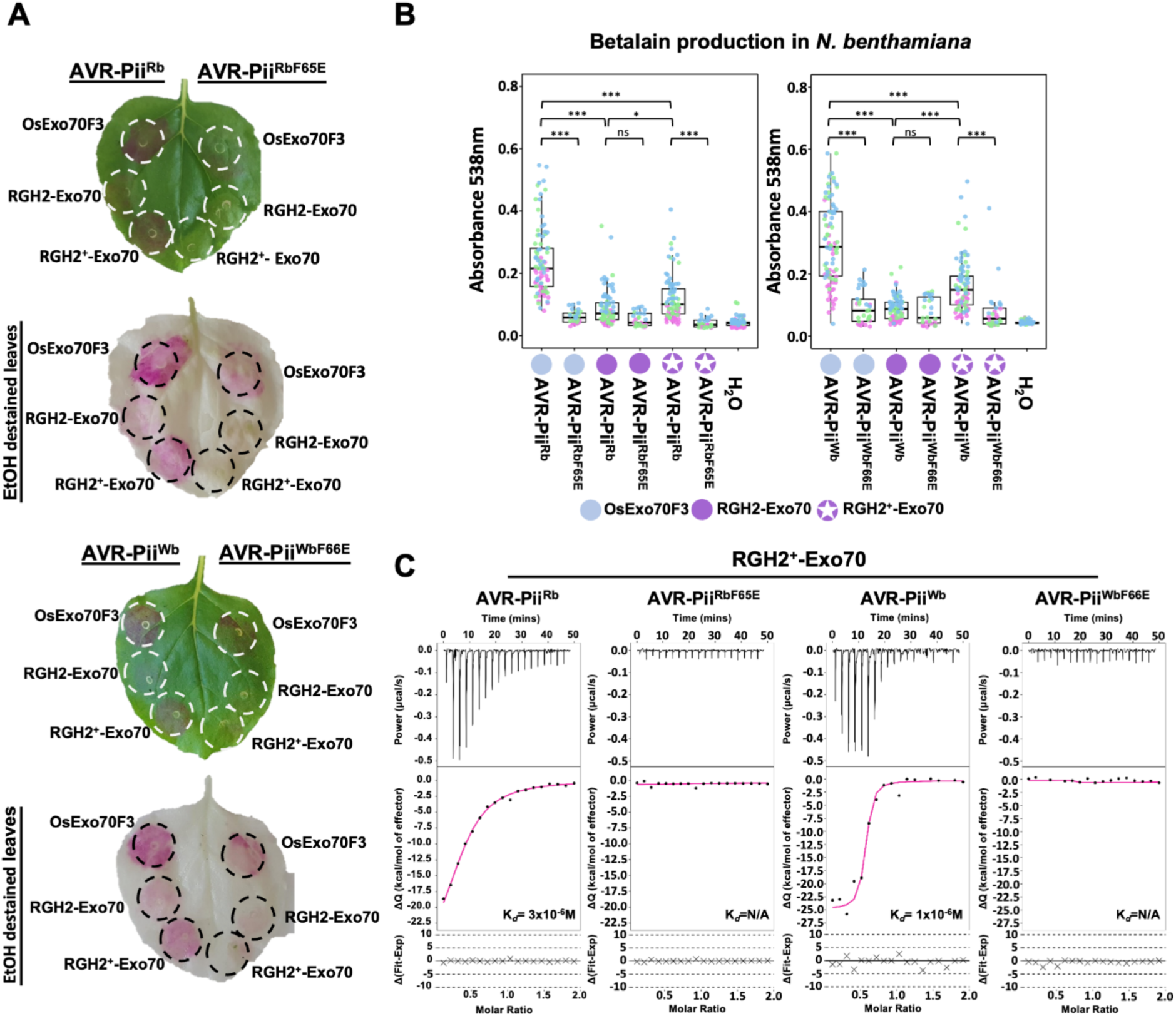
AVR-Pii^Wb^ exhibits increased binding to RGH2^+^ compared with RGH2 in planta and in vitro. **A.** Betalain production in *N. benthamiana* leaves following Agrobacterium infiltration with OsExo70F3-GAL4-BD, RGH2-Exo70-GAL4-BD or RGH2^+^-Exo70-GAL4-BD and rice or wheat blast VP16-GAL4-AD-AVR-Pii variants or mutant effectors. Leaves expressing RGH2^+^ show increased betalain accumulation compared to RGH2 with AVR-Pii variants. Upper panels show brightfield images of *N. benthamiana* leaves three days post infiltration. Lower panel shows the same leaves cleared overnight in 100% ethanol. Infiltration points are marked with a white or black circle. **B.** Leaves cleared with ethanol were used for betalain extraction. The absorbance for betalain production was measured at OD538nm from independent infiltration sites for each construct combination. Replicates are designated with different colours (magenta-Replicate 1 (R1), green-Replicate 2 (R2), blue-Replicate 3 (R3)). Box plots represent the median (horizontal line), upper and lower quartiles (boxes), and 1.5× interquartile range (whiskers). Asterisks indicate statistically significant differences from the control (NS: not significant, \**p* < 0.05, \*\**p* < 0.01, \*\*\**p* < 0.001, one-way ANOVA followed by Tukey’s test). Data shown are from a pool of three biological replicates, n = 15 plants per pool (30 infiltration points per one biological replicate). **C.** Binding of rice blast AVR-Pii, the wheat blast variant, and mutants to RGH2^+^-Exo70 in vitro, as determined by ITC. Upper panel represents the heat differences upon injection of effector variants and mutants into the cell containing RGH2^+^-Exo70. Middle panel shows the integrated heats of injection (dots - black colour) and best fit (solid line - magenta colour) to a single-site binding model calculated using AFFINImeter ITC analysis software^68^. Bottom panel indicates the difference between the fit to a single- site binding model and the experimental data. The experiments shown are representative of three replicates.

### RGH2^+^-Exo70 binds AVR-Pii^Rb^ and AVR-Pii^Wb^ more tightly compared to wild-type in vitro

To extend the in vivo analysis, we purified RGH2^+^-Exo70 from *E. coli* and tested binding to AVR-Pii^Rb^ and AVR-Pii^Wb^ by ITC. Consistent with the RUBY assay, the binding affinity of RGH2^+^-Exo70 to AVR-Pii^Rb^ and AVR-Pii^Wb^ was increased compared to RGH2-Exo70, with a dissociation constant (*KD*) of 3.12×10^−6^ M and 1.06×10^−6^ M, respectively (Fig. 4C). No binding to RGH2^+^-Exo70 was observed with AVR-Pii^RbF65E^ or AVR-Pii^WbF66E^ (Fig. 4C). These in vitro experiments confirmed tighter binding of RGH2^+^-Exo70 towards AVR-Pii^Rb^ and AVR-Pii^Wb^ compared to the wild type RGH2-Exo70.

### RGH2^+^/RGH3 enhanced cell death responses to AVR-Pii^Rb^ and AVR-Pii^Wb^ in *N. benthamiana*

We hypothesised that increased effector binding to RGH2^+^-Exo70 may translate into improved immune responses in plants compared to RGH2. To test this, we first co-expressed RGH2 and RGH3 in *N. benthamiana* alone to assess potential autoactivity, and with AVR-Pii^Rb^ or AVR-Pii^Wb^ to observe effector-dependent cell death. Imaging leaves three days post infiltration revealed that no macroscopic cell death was observed for RGH2/RGH3 alone, or when co-expressed with GFP or with effector Phe to Glu mutants (Fig. 5A, 5B). However, cell death was observed under both white light and UV light on co-expression of RGH2/RGH3 and either of AVR-Pii^Rb^ or AVR-Pii^Wb^ (Fig. 5A, 5B). In parallel, to test the impact of RGH2^+^ on cell death, we co-expressed this NLR with RGH3, both with and without effectors. Again, we did not observe autoactivity, or cell death with negative controls. Intriguingly, we observed qualitatively stronger cell death in leaf spots infiltrated with RGH2^+^ compared to RGH2, and the cell death was stronger with AVR-Pii^Wb^ over AVR-Pii^Rb^ (Fig. 5A, 5B). To quantify the cell death on expression of NLRs and effectors in *N. benthamiana*, we measured electrolyte leakage from leaf disks at 4, 24, and 48 hours post-infiltration (Fig. 5C, S12A, B). Here, we observed significant increases in conductivity from samples of RGH2^+^/RGH3-infiltrated tissue with AVR-Pii^Wb^ over AVR-Pii^Rb^ compared to RGH2/RGH3. RGH2/RGH3 or RGH2^+^/RGH3 with AVR-Pii^F65E^ displayed minimal change in conductivity compared with RGH2/RGH3 and RGH2^+^/RGH3 alone (Fig. 5C, Supplementary Fig. S12A, B). We also observed higher levels of conductivity in samples with AVR-Pii^Wb^ compared to AVR-Pii^Rb^, in agreement with the macroscopic cell death observations. Protein accumulation was assessed by western blot for all *N. benthamiana* cell death experiments (Fig. 5D, E). Collectively, these data demonstrate that RGH2^+^/RGH3 exhibits a stronger cell death response towards the effectors compared to RGH2/RGH3 in *N. benthamiana*, and that AVR-Pii^Wb^ elicits a stronger response than AVR-Pii^Rb^.

**Fig. 5:**
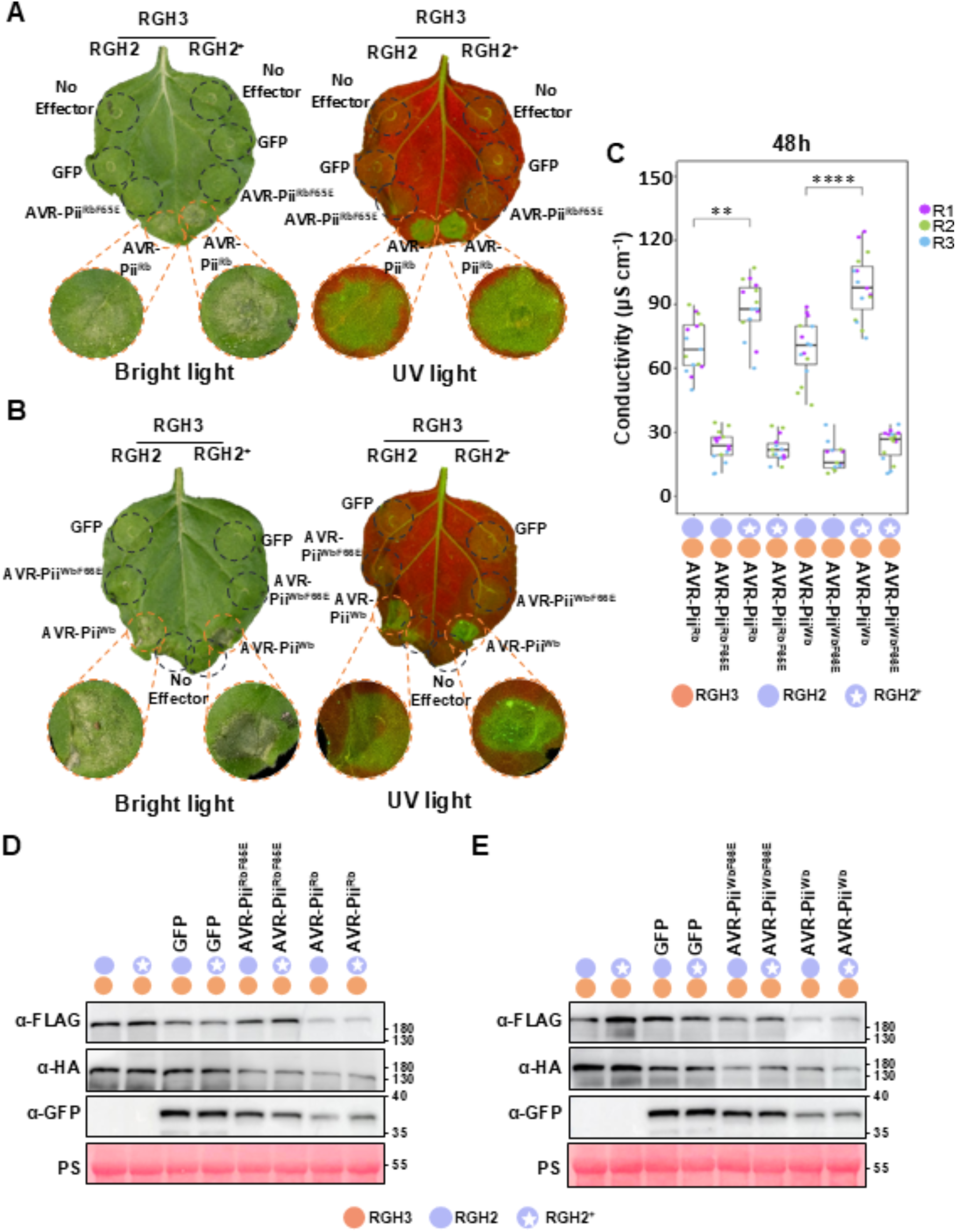
RGH2^+^/RGH3 elicit an increased cell death response to rice blast AVR-Pii and the wheat blast variant compared to RGH2/RGH3. **A.** Macroscopic cell death phenotypes observed on transient expression of RGH2/RGH3 or RGH2^+^/RGH3 NLR pairs with GFP, GFP-AVR-Pii^Rb^ or GFP-AVR-Pii^RbF65E^ mutant in *N. benthamiana*. Leaves were photographed five days post infiltration and are representative of three independent experiments. **B.** As for Fig. 5A but with the AVR-Pii^Wb^ variant and AVR-Pii^WbF66E^ mutant. **C.** Conductivity assay from *N. benthamiana* tissue infiltrated with RGH2/RGH3 or RGH2^+^/RGH3 NLR pairs with rice or wheat blast GFP-AVR-Pii variants or mutants at 48 hours post infiltration (hpi). Measurements of ion leakage after 4 and 24 hpi are shown in Supplementary Fig.12. Box plots represent the median (horizontal line), upper and lower quartiles (boxes), and 1.5× interquartile range (whiskers). Statistically significant differences are denoted by an asterisk (NS: not significant, \*\**p* <0.01, \*\*\*\**p* <0.0001 Student’s t-test). Data from three independent experimental replicates are shown (n=5 plants per experiment). Replicates are designated with different colours (magenta-Replicate 1 (R1), green-Replicate 2 (R2), blue-Replicate 3 (R3)). **D.** Expression of constructs used for the cell death and conductivity assays in *N. benthamiama* probed by western blot. RGH2-Exo70-FLAG, RGH2^+^-Exo70-FLAG, RGH3-HA, the rice blast effector and AVR-Pii^RbF65E^ mutant were detected with the appropriate antibodies as labelled. Ponceau S staining shows the amount of total protein loaded per lane in the input. **E.** As for Fig. 5D but with the AVR-Pii^Wb^ variant and AVR-Pii^WbF66E^ mutant.

### Enhanced recognition of AVR-Pii^Wb^ by RGH2^+^/RGH3 in transgenic barley plants

We hypothesised that the increase in cell death response of RGH2^+^/RGH3 to effectors compared with RGH2/RGH3 in *N. benthamiana*, could confer improved disease resistance in barley to pathogens encoding these effectors. For this, we focussed on AVR-Pii^Wb^ due to the need to explore novel resistances against wheat blast^52^.

The wild-type RGH2/RGH3 was previously cloned with the native promoter and terminator and transformed into the barley cultivar Golden Promise^46^. The RGH2/RGH3 genomic transformation construct was edited using site-directed mutagenesis to covert the integrated Exo70 binding site into RGH2+ and was transformed into the barley cultivar Golden Promise using Agrobacterium-mediated transformation.

RGH2^+^/RGH3 and RGH2/RGH3 transgenic T1 families were tested for recognition of AVR-Pii^Wb^ by inoculation with *M. oryzae* isolate Guy11 (lacking AVR-Pii) transformed with AVR-Pii^Wb^ (Golden Promise is susceptible to Guy11). Spray inoculation on detached leaves was used to assess phenotypes, which were scored on a macroscopic 0-4 scale for disease severity^53^(Fig. 6A, C). For RGH2/RGH3 transgenics, four independent T1 families were screened with 84, 72, and 82 individuals screened across experimental replicates. For RGH2^+^/RGH3 transgenics, four independent T1 families were screened with 119, 91, and 93 individuals across replicates. A total of 88, 117, and 131 individuals of the susceptible Golden Promise transgenic background were screened across replicates (Figure 6B).

**Fig. 6:**
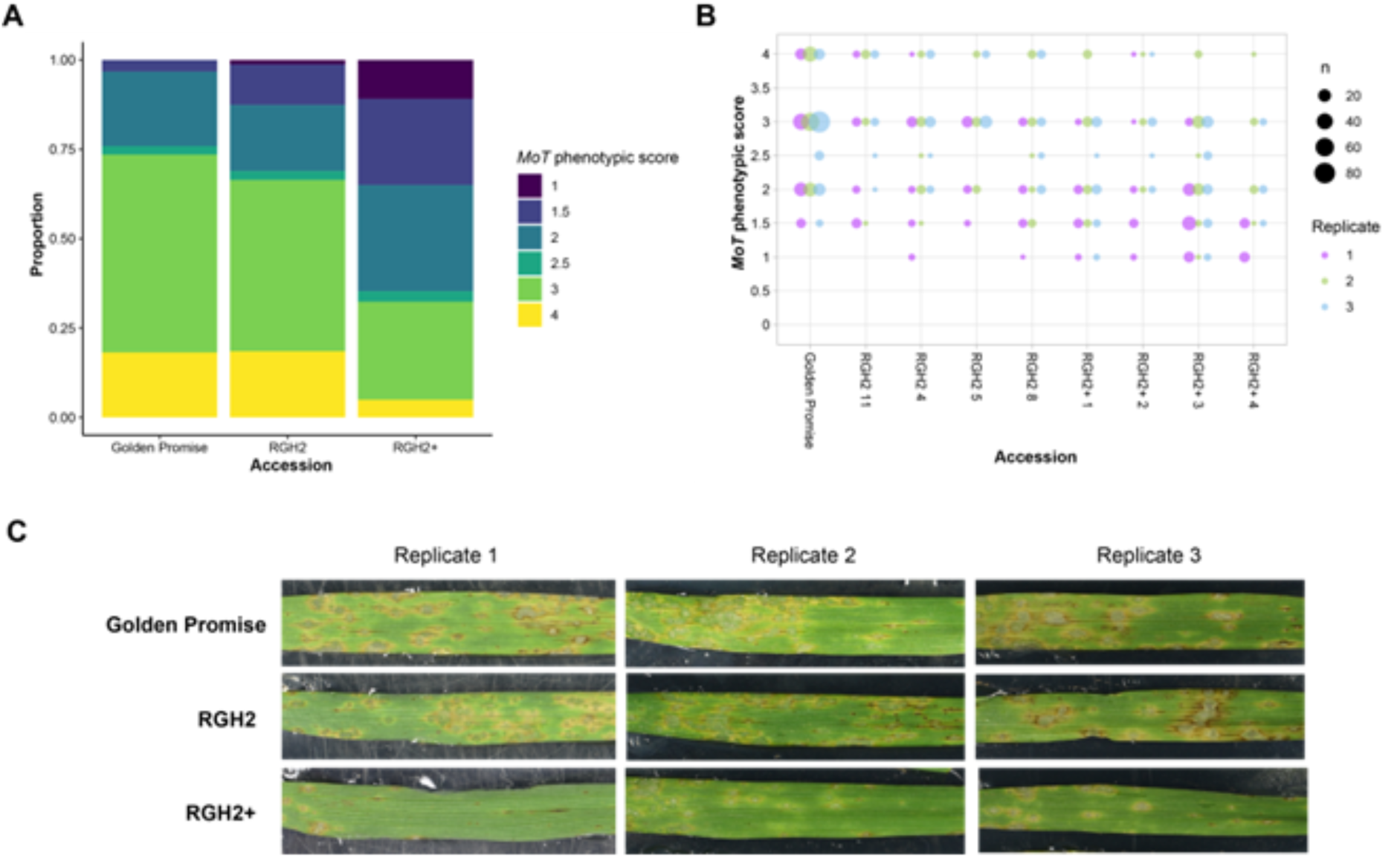
RGH2^+^/RGH3 transgenic lines show reduced disease severity to *M. oryzae* isolates carrying AVR-Pii^Wb^. **A.** Proportional phenotypes of individuals across Golden Promise control, RGH2, and RGH2+ transgenic lines. Colours indicate ordinal phenotype classes on the 0 – 4 phenotype scale from resistant (0) to susceptible (4). **B.** Phenotypic scores from individuals in independent T1 family of RGH2/RGH3 and RGH2+/RGH3 transgenic lines and the Golden Promise control. Phenotypes plotted on the 0 – 4 phenotype scale from resistant (0) to susceptible (4). Circle size indicates the number of individuals with each phenotypic score for each replicate. **C.** Macroscopic images of spray inoculations on detached leaves of barley accessions infected with *M. oryzae* isolate Guy11 transformed with AVR-Pii^Wb^. Representative phenotypes of individuals from T1 families of RGH2/RGH3 and RGH2^+^/RGH3 transgenics and Golden Promise (the transgenic background) are shown from each experimental replicate.

To evaluate the effect of RGH2^+^/RGH3 and RGH2/RGH3 on disease severity, we used a cumulative link mixed model fitted with the Laplace approximation to assess the ordinal infection scores. The model included the accession (RGH2^+^/RGH3, RGH2/RGH3, and Golden Promise) as a fixed effect, with random effects of experimental replicate, independent transgenic families, and the interaction between replicate and transgenic family. For the random effects, variability is observed across replicates (σ² = 0.83, SD = 0.91), independent families perform slightly differently across replicates (σ² = 0.37, SD = 0.61), and plants within the same independent family show minimal variation (σ² = 0.01, SD = 0.10). A significant difference was observed between RGH2^+^/RGH3 transgenics compared to the Golden Promise control (β = -1.90, SE = 0.44, z = -4.28, p < 0.0001). By contrast, RGH2/RGH3 transgenic lines showed no significant difference from the control (β = -0.20, SE = 0.44, z = -0.46, p = 0.648), suggesting that RGH2/RGH3 alone does not confer reduced disease severity under our experimental conditions (Fig. 6A, B).

Further, post-hoc pairwise comparisons of estimated marginal means using Tukey’s HSD confirmed these results. RGH2^+^/RGH3 transgenic lines demonstrated significantly reduced disease severity to Guy11 transformed with AVR-Pii^Wb^ compared to both RGH2/RGH3 transgenic lines (difference = 1.70, SE = 0.32, p < 0.001) and Golden Promise (difference = 1.90, SE = 0.44, p < 0.001). No significant difference was observed between RGH2/RGH3 and Golden Promise (difference = 0.20, SE = 0.44, p = 0.891) (Fig. 6A, B). Therefore, engineered RGH2^+^/RGH3 confers reduced disease severity compared to wild-type RGH2/RGH3.

## Discussion

NLR immune receptors play a crucial role in protecting plants against pathogens and are a critical component of disease resistance breeding programs in crops. The diverse effector repertoires of pathogens and their ability to adapt to immune systems presents challenges that can limit the effectiveness of NLRs deployed in agriculture. Approaches to address this limitation include engineering NLRs to broaden their recognition spectrum to target conserved effectors across pathogen isolates. Here, we have introduced structure-guided mutations in the effector binding interface of the Exo70 integrated domain of the barley NLR immune receptor RGH2 to enhance binding to rice and wheat blast effectors. Using a combination of in vitro and in vivo experiments, we demonstrate this engineered RGH2^+^ shows higher affinity binding towards rice blast AVR-Pii and the wheat blast variant compared to RGH2, leading to improved immune responses and reduced pathogen virulence in transgenic barley lines.

With the emergence of new disease pandemics, such as wheat blast that threatens yields across Asia and Africa, there is a pressing need for identifying new sources of resistance^54–56^. Screening wild species and landraces for novel resistance traits continues to deliver new genes and germplasm for breeding^57^. Alternative approaches are enabled by advances in understanding the molecular mechanisms of plant immune receptor structure and function. Such knowledge can be deployed through engineering and improved transformation pipelines, alongside ongoing improvements in gene-editing tools.

NLR integrated domains (NLR-IDs) bait pathogen effectors for immune activation in cereals and other plants^14,58–60^. They present opportunities for receptor engineering, to optimise effector recognition ahead of activating conserved downstream mechanisms of NLR-mediated immunity^14,59^. Exo70 integrated domains are under-studied compared to other examples such as HMA and WRKY domains^13,14,16,20,59,60^. The best studied example of NLR-mediated immunity requiring Exo70 proteins is towards the *M. oryzae* effector AVR-Pii. However, in this case the Exo70 protein is guarded by the NLR pair Pii, and not integrated within an immune receptor^30,31^. In plants, Exo70 proteins form an expanded family with increased gene copy number and functional diversification^35,36,38,43^ and they are targeted by effectors from diverse pathogens^44^. Our study highlights the potential of engineering NLR integrated Exo70 domains to mediate activation of immunity.

Our rationale for RGH2-Exo70 engineering was guided by the interface formed in the crystal structure of the OsExo70F2/AVR-Pii complex^29^. The mutations to generate RGH2^+^ resulted in subtle changes to amino acid properties at the interface, including Alanine to Valine (position 1378), Isoleucine to Leucine (position 1379) and Leucine to Methionine (position 1396). We hypothesised these differences could explain the reduced affinities of AVR-Pii^Rb^ and AVR-Pii^Wb^ to RGH2-Exo70 over OsExo70F3, and ultimately improve the binding and response of the RGH2^+^/RGH3 pair. Subtle changes acquired in effectors through evolution can result in escape from detection by plant immune receptors, potentially reducing their effectiveness in crops. By contrast, we show here that subtle amino acid changes can be harnessed to engineer receptors to boost plant immunity, including to effector families conserved across pathogen isolates. These changes maybe more broadly suitable for targeted gene-editing approaches in agriculture, balanced against larger scale genetic modifications (e.g. domain swapping or full gene transformation). However, while OsExo70F3 binds AVR-Pii^Rb^ with nanomolar affinity, RGH2^+^ binding to AVR-Pii^Rb^ and AVR-Pii^Wb^ remains within the micromolar range. This suggests there may be opportunities, beyond the scope of this study, to further improve effector binding to RGH2 and response to wheat blast expressing AVR-Pii^Wb^.

Effectors form diverse sequence and structural repertoires across different pathogen isolates^61–63^. AVR-Pii^Rb^ and AVR-Pii^Wb^ are both members of tribe VIII of the Zinc-finger (ZiF) effector family^32^, sharing 72 % sequence identity. We observed that AVR-Pii^Rb^ and AVR-Pii^Wb^ elicit different levels of RGH2/RGH3 activation, which was even clearer with RGH2^+^/RGH3. Specifically, AVR-Pii^Wb^ elicited stronger cell death compared to AVR-Pii^Rb^ in *N. benthamiana* assays (Fig. 5A). ZiF effectors from different tribes (including tribe VIII) are prevalent in wheat blast lineages compared to rice blast, where many are either absent or are pseudogenes^32^. While we focussed on engineering the barley NLR RGH2 receptor pair towards AVR-Pii^Rb^ and AVR-Pii^Wb^ here, there are future opportunities to expand responses towards other ZiF family members by exploring surface complementarity between predicted effector structures and the binding sites of Exo70 integrated domains using computational approaches^64^.

In particular, this includes members from ZiF effector Tribes VII and IX, as these were previously shown to bind OsExo70F3^32^.

In this study, we aimed to engineer new resistance to effectors from the blast pathogen (specifically wheat blast) and in general broaden our understanding of NLR-mediated resistance mechanisms. Engineering of the NLR RGH2, guided by the binding of rice blast AVR-Pii to OsExo70F2, reduced the disease severity of *M. oryzae* when introduced in transgenic barley. Additional work is required to explore the potential to increase resistance further, possibly through new enhanced receptor-effector binding variants or optimising NLR expression levels.

### Conclusion and outlook

Engineering NLR integrated domains is a promising path to expand the effector recognition profile of these plant immune receptors. Previously, the rice NLR pairs Pik (Pik-1/Pik-2) and Pia (RGA4/RGA5), which both contain integrated HMA domains in their sensor NLRs to bait effectors, have been engineered with novel response profiles ^19,20,28,65^. Here, we further the concept of NLR integrated domain engineering to the Exo70 domain of the RGH2/RGH3 pair, demonstrating its promise as a scaffold to expand the spectrum of effector recognition specificities.

Looking ahead, future directions could involve applying this system to wheat cultivars. Interestingly, RGH2/RGH3 is present in rice, however RGH2 has an integrated thioredoxin domain rather the Exo70^47^. As AVR-Pii has been lost from most *M. oryzae* isolates infecting rice (likely due to deployment of Pii-based resistance in rice) the evolution of different integrated domains in the RGH2/RGH3 architecture supports this chassis as an appropriate target for engineering. Finally, it may be possible to engineer resistance to other pathogens by installing additional diverse Exo70 domains, or other unrelated effector targets.

## Method details

### Plasmids and cloning procedures

*Escherichia coli* Stellar™ (Takara Bio), SHuffle (Invitrogen) and Rosetta™ (DE3) cells were used for cloning and protein expression. All plasmids were generated by either In-Fusion^®^ cloning (Takara Bio, USA) or the Goldengate system^66^ using resources from TSL Synbio. All plasmids are described in **Supplementary Table 1**, enabling full details of constructs used, including epitope tags. The modules used were either amplified by PCR, synthesized as PCR products (Gblocks, GeneWiz) or obtained from TSL SynBio. Restriction enzymes were purchased from New England Biolabs (Ipswich, MA, USA). Primers and their purpose are listed in **Supplementary Table 2**.

### Transcriptome assembly and phylogenetic analysis

Publicly available RNAseq data sets for Poales species were identified from NCBI SRA database and *de novo* transcriptome assemblies were generated using Trinity with default parameters (version 2013-11-10). The Poales species phylogenetic tree was generated using the QKbusco pipeline (https://github.com/matthewmoscou/QKbusco) as described in Brabham *et al.,* (2025)^53^. Briefly, open reading frames from assemblies predicted using TransDecoder (v4.1.0). Benchmarking universal single-copy orthologs were identified using BUSCO (v3.0.2) with default parameters and embryophyte_odb9 library. Genes identified with BUSCO were used to generate a multiple sequence alignment using QKbusco_merge.py using parameter ‘allfragmented’. Codon-based multiple sequence alignment was performed with PRANK (v.170427) using default parameters. Multiple sequence alignments were merged using QKbusco_phylogeny.py with coverage depth of 40% at individual sites. The maximum likelihood phylogenetic tree was generated using RAxML (v8.2.12) with the GTRGAMMA model.

InterProScan (v5.27-66.0) using default parameters was used to identify NB-ARC (Pfam PF00027 and PF00931) and Exo70 (Pfam PF03081) domain containing proteins from *de novo* transcriptomes predicted proteins from TransDecoder. Domain boundaries of integrated NLRs were manually curated based on Pfam annotation. Phylogenetic analysis of Exo70 domains was carried out as described by Holden *et al.*, (2022)^43^. Briefly, structure-guided multiple sequence alignment was performed with MAFFT software (v7.481) using DASH (Database of Aligned Structural Homologs) with default parameters. Structurally resolved Exo70 structures included *A. thaliana* Exo70A1 (PDB 4RL5), *S. cerevisiae* (yeast) Exo70 (PDB 2B1E and 5YFP), and *Mus musculus* (mouse) (PDB 2PFT). The maximum likelihood phylogenetic tree was constructed using RAxML (v8.2.12) with the JTT amino acid substitution model, gamma model of rate heterogeneity, and 1,000 bootstraps. iTOL (Interactive Tree of Life) was used for phylogenetic tree visualization, and *S. cerevisiae* and *M. musculus* Exo70 were used as outgroups.

### Cell death assay

Four-week-old *N. benthamiana* plants were simultaneously infiltrated with Agrobacteria carrying the relevant constructs *35Spro::*RGH2-FLAG, *35Spro::*RGH2^+^-FLAG, *35Spro::*RGH3-HA and *MASpro::*GFP-AVR-Pii from wheat and rice or *35Spro::*GFP-HA and *MASpro::*GFP-AVR-Pii^F65E^ as controls. The Agrobacterium strains were resuspended in ARM buffer (Agrobacterium resuspension medium) containing 10 mM MES (pH 5.6), 10 mM MgCl2, and 150 μM acetosyringone. The optical density at 600 nm (OD600) was adjusted as follows: RGH2/RGH2^+^ and RGH3 at 0.4, effectors and controls at 0.6, and p19 at 0.1. Two leaves from each *N. benthamiana* plant were infiltrated using a needleless syringe. Leaves were harvested at 3 days post-infiltration (dpi), and UV fluorescence images were captured from the abaxial side. Imaging was conducted using a Nikon D4 camera equipped with a 60-mm macro lens, with ISO set at 1600 and an exposure time of approximately 10 seconds at F14. A Kodak Wratten No.8 filter was used, and the white balance was set to 6,250 °K. Illumination was provided by Blak–Ray longwave (365 nm) B-100AP spotlight lamps, which were moved around the subject during exposure to ensure even illumination. The presented images are representative of three independent experiments, each with a minimum of 6 leaves analysed.

### Ion leakage assays

Leaves of *N. benthamiana* were infiltrated with either *35Spro::*RGH2-FLAG or *35Spro::*RGH2^+^-FLAG in combination with *MASpro::*RGH3-HA and either *MASpro::*GFP-AVR-Pii or *MASpro::*GFP-AVR-Pii^Phe/Glu^ from both rice and wheat blast. Ion leakage assays were performed as described in Hatsugai and Katagiri (2018)^67^. Briefly, measurements were performed after 4, 24 and 48 hours post-harvest with a compact conductivity meter (LAQUAtwin-EC-33, Horiba Scientific). Five leaf discs (4mm diameter) were harvested from different leaves and immersed inside 2ml ddH2O as one biological replicate. Each measurement contains five biological replicates, and all experiments were performed at least three times.

### Split GAL4 RUBY assay

To detect protein-protein binding in *N. benthamiana* we performed the split GAL4 RUBY assay according to Chen *et al*., 2023.^51^ Briefly, Agrobacterium containing OsExo70F3, RGH2-Exo70 and RGH2^+^-Exo70 (residues 1033-1637) fused to the yeast GAL4 DNA binding domain protein under the *35S* promoter. Wheat and rice blast AVR-Pii and the mutants or ZiF_IIb were fused to the herpes virus VP16 AD either under the *35S* promoter or *MAS* promoter. *N. benthamiana* leaves were harvested after 3dpi. To extract betalain, harvested plant leaves were completely immersed in 100% ethanol until all chlorophyll was removed. A diameter of 0.8 cm was taken from each infiltration site and put into H20. Extracted betalain accumulation was measured in a Greiner F-bottom clear plate in a microplate reader (Clariostar, BMG Labtech) at 538 nm wavelength. All experiments were performed with five plants and repeated at least 3 times.

To confirm protein expression, plant samples were harvested 3 dpi, ground in liquid nitrogen and heated for 10 min at 70 °C in LDS buffer (Bio-Rad Laboratories, USA). Samples were centrifugated and the supernatant was subjected to SDS-PAGE gels and western blot. The membranes were probed with anti-GAL4 DNA-BD_HRP (Santa Cruz Biotechnology) for OsExo70F3, RGH2-Exo70 and RGH2^+^-Exo70 and with the anti-VP16 antibody (Sigma-Aldrich, USA, V4388) primary antibody and Goat anti-rabbit as secondary antibody (Sigma-Aldrich, USA, 12-348) for the effectors.

### Co-immunoprecipitation

Transient gene expression in planta was performed by co-infiltrating 4-week-old *N. benthamiana* plants with *A. tumefaciens* strain GV3101 carrying *35Spro::*OsExo70F3-FLAG or *35Spro::*RGH2-Exo70-FLAG with rice AVR-Pii, the wheat blast variant or their respective mutants. NLRs *MASpro::*FLAG-RGH2, *MASpro::*FLAG-RGH2^+^, *MASpro::*HA-RGH3 and rice AVR-Pii, the wheat blast variant, or their respective mutants, and ZiF_IIb under the *MAS* promoter (Fig. 3D and Fig. S9) were infiltrated at OD600 0.4 and 0.6, respectively, in ARM buffer (Agrobacterium resuspension medium, 10 mM MgCl2, 10 mM 2-(N-morpholine)-ethanesulfonic acid [MES], and pH 5.6) with the addition of 150 µM acetosyringone. Two leaves from each *N. benthamiana* plant were infiltrated and 48 h later, samples were harvested and frozen in liquid nitrogen. Total proteins were extracted from 450 mg tissue samples and suspended in 2 mL GTEN buffer (25 mM Tris pH 7.5, 150 mM NaCl, 1 mM EDTA, 10% v/v glycerol, 2% w/v PVPP, 10 mM DTT, 1 × complete protease inhibitor tablet per 50 mL (Roche), and 0.1% Tween 20). Extracts were cleared by centrifugation at 4000 xg at 4 °C for 30 min. Protein pull downs were performed by adding 30 µL of anti-FLAG magnetic beads using M2 anti-FLAG magnetic beads (Sigma). Briefly, samples were incubated in a rotary mixer for 1h at 4 °C, washed afterwards 5 times with 1 ml IP buffer (25 mM Tris pH 7.5, 150 mM NaCl, 1 mM EDTA, 10% glycerol, and 0.1% Tween 20) and proteins were eluted by incubating the beads at 70°C in 30 µL of LDS RunBlue sample buffer for 10 mins. About 15–20 μL of the extracts were analysed by SDS-PAGE followed by immunoblot using either anti-FLAG (Cohesion Biosciences, at 1:3000 dilution), anti-GFP (Santa Cruz Biotechnology, at 1:3000 dilution) or HA-probe (Santa Cruz Biotechnology, 1:5000 dilution). Experiments were repeated at least three times.

### Protein immunoblotting

Protein samples were prepared from five tissue discs (8 mm diameter) sampled from *N. benthamiana* leaves at 2 or 3 days after agroinfiltration and were homogenized in 100 μL of 2X LDS loading buffer (Bio-Rad). Proteins were transferred onto a PVDF (polyvinylidene difluoride) membrane using Trans-Blot Turbo transfer system (Bio-Rad) according to the manufacturer’s protocol. Immunoblotting was performed with HA-probe (Santa Cruz Biotechnology, 1:5000 dilution), anti-GFP antibody ((Santa Cruz Biotechnology, at 1:3000 dilution) or anti-FLAG (Cohesion Biosciences, at 1:3000 dilution). Proteins were visualized using Clarity Max Western ECL Substrate (Bio Rad) in the ImageQuant LAS 500 spectrophotometer (GE Healthcare). Total protein loading was visualized by staining the PVDF membrane with Ponceau S solution (Sigma-Aldrich, P7170).

### Yeast two-hybrid assay

To examine the interaction between all Exo70s and effectors we used the Matchmaker Gold system (Takara Bio, USA) following the manufacturer’s protocol. The proteins were cloned into pGBKT7 or pGADT7 vectors, respectively. Exo70 integrated domains RGH2-Exo70 (residues 1033-1637), RGH2^+^-Exo70 (residues 1033-1637), Apu28443-Exo70 (residues 990-1644), Hla80948-Exo70 (residues 765-1269), Hvu23924-Exo70 (residues 726-1321), Jef9120-Exo70 (residues 364-887) were cloned into pGBKT7 for GAL4 binding domain fusion. OsExo70F3 in pGBKT7 and ZiF effectors and mutants in pGADT7 were previously generated^32^. Constructs were co-transformed into chemically competent Y2HGold cells using a Frozen-EZ Yeast transformation II Kit (Zymo research, USA). We selected yeast co-transformants on selective dropout (SD) media lacking only Trp and Leu. Protein interactions were assessed on SD selection medium lacking Trp, Leu, His and Ade containing X-α-Gal and supplemented with 0.2 ug/ml Aureobasidin A (Takara Bio, USA). Plates were incubated on 30 °C and interaction was observed 4 days later. Each experiment was repeated a minimum of three times with similar results.

To confirm protein expression, total yeast extracts from transformed colonies were produced by heating the cells for 10 min at 70 °C in LDS buffer (Bio-Rad Laboratories, USA). Samples were centrifugated and the supernatant was subjected to SDS-PAGE gels and western blot. The membranes were probed with anti-GAL4 DNA-BD (Sigma Aldrich, USA, G3042) for OsExo70F3, RGH2-Exo70, RGH2^+^-Exo70, Apu28443-Exo70, Hla80948-Exo70, Hvu23924-Exo70, Jef9120-Exo70 proteins in pGBKT7 and with the anti-GAL4 activation domain (Sigma Aldrich, USA, G9293) antibodies for effectors in pGADT7.

### Protein expression and purification

RGH2-Exo70 and RGH2^+^-Exo70 domains (residues 1033-1637) were 6xHis-SUMO-tagged and produced in *E. coli* Rosetta™ (DE3) and AVR-Pii variants were 6xHis-MBP-tagged and produced in *E. coli* SHuffle cells using established protocols^29^. All tags were added at the N-terminus. In brief, cell cultures were grown in LB media at 37 °C for Rosetta and 30 °C for SHuffle cells for 5–7 hr and then at 16 °C overnight. Cells were harvested by centrifugation (10 min, 5500x g, 4 °C) and re-suspended in 50 mM Tris-HCl (pH 7.5), 500 mM NaCl, 50 mM glycine, 5% (vol/vol) glycerol, and 20 mM imidazole supplemented with cOmplete EDTA-free Protease Inhibitor Cocktail (Roche). Cells were lysed by sonication and clarified by centrifugation (50,000x g for 30 mins at 4 °C). Protein purification was carried out as previously described^29^.

### Isothermal titration calorimetry

ITC experiments were performed using a MicroCal PEAQ-ITC instrument (Malvern, United Kingdom). For each experiment, 300 μL of RGH2-Exo70 or RGH2^+^-Exo70 domains at a concentration of 20 μM was loaded into the calorimetric cell, while the syringe contained 200 μM of either wheat or rice blast AVR-Pii or their Phe to Glu mutants. Each run consisted of a single injection of 0.5 μL followed by 18 injections of 2 μL each, with intervals of 120 s between injections. The stirring speed was 750 rpm. Data obtained from the experiments were analysed using AFFINImeter ITC software^68^. Both wild type and mutant samples were subjected to triplicate runs. The experiments were conducted at 25 °C in a buffer containing 20 mM HEPES (pH 7.5), 150 mM NaCl, and 5% (vol/vol) glycerol. The stability of RGH2/RGH2^+^ Exo70 domains on purification likely affects the accurate measurement of stoichiometry on complex formation with the effectors (1:1 expected based on the structure of OsExo70F2 with AVR-Pii).

### *M. oryzae* inoculation and plant phenotypic assays

Protocols for culturing and inoculation of *M. oryzae* were similar to those previously described^46,69,70^. Briefly, hyphal tips from culture plates were transferred to oatmeal agar (20 g oatmeal, 10 g agar, 2.5 g sucrose, and addition of ddH2O to 500 mL) plates (deep petri dish 100 × 20 mm) and incubated for 10 to 15 d at 24 °C to produce conidia. Oatmeal plates were washed with 8 mL dH2O and conidia collected by gentle scraping with the tip of a 1.5 mL tube. The suspension was filtered through Miracloth (Merck Chemicals, Ref.: 475855-1r) and collected. Spore concentration was counted with a hemocytometer and adjusted to 1 × 10^5^ spores per milliliter in dH2O with 0.01% Tween 20 (Merck Chemicals, CAS Number: 9005-64-5).

For detached leaf assays, barley leaves were sampled and placed on agar in boxes (2.5 g agar-agar [Fisher, CAS 9002-18-0]; 50 mL benzimidazole [1 g/1 L H2O stock solution]; 450 mL H2OFor spray inoculations, the conidial suspension was sprayed onto the leaves in each box using a small atomiser spray bottle with 3 - 4 sprays used until droplets were observed on the leaf surface. Boxes were placed in a Sanyo cabinet and maintained at 25 °C. Continuous light was used in the first 24 hrs and then, after excess water was removed from droplets using sterile Miracloth, boxes were maintained in a 16/8 hr light/dark cycle. Phenotypes were recorded at 7 dpi using a macroscopic phenotype scale^46^ (0 = complete resistance; 1 = small brown resistant spots; 2 = susceptible larger eyespot lesions; 3 = larger spreading lesions; and 4 = hypersusceptibility).

### Statistical analyses

Graphs and statistical analysis (the Student’s *t* test, Tukey’s ANOVA, Two-Way ANOVA) were performed using GraphPad Prism version 10.2.2 and R-Studio version 2024.04.2+764. Ordinal phenotyping data were analysed using a cumulative link mixed model implemented in the R package ordinal (version 2023.12.4.1; https://CRAN.R-project.org/package=ordinal)^71^. The model was specified as: phenotype ∼ accession + (1 | independent family) + (1 | replicate) + (1 | replicate:independent family) where accession specifies Golden Promise control, RGH2, or RGH2+ transgenic lines. The Laplace approximation was used for likelihood estimation. Post hoc pairwise comparisons were conducted using the estimated marginal means derived using the emmeans package (version 1.11.2.8; https://CRAN.R-project.org/package=emmeans) and *p*-values were adjusted using Tukey’s method. Details of plots and statistical tests used are given in the appropriate figure legends.

## Data/Resource availability

Further information and requests for resources and reagents should be directed to the Mark Banfield. There are no restrictions on the availability of materials. Any additional information required to re-analyze the reported data is available from the lead contact. The data, including FASTA, Phylip, and Nexus files used for preparing Fig. 1 are available on Figshare (www.doi.org/10.6084/m9.figshare.28057085). All other study data are included in the article and/or Supplementary Information.

## Acknowledgments

The authors thank the Biotechnology and Biological Sciences Research Council (UKRI-BBSRC, UK, grants BB/V015508/1, BB/P012574, BBS/E/J/000PR9795, BBS/E/J/000PR9796 and BB/X010996/1), the BBSRC Norwich Research Park Biosciences Doctoral Training Partnership (grant BB/T008717/1), the John Innes Foundation, the Gatsby Charitable Foundation, the United States Department of Agriculture-Agricultural Research Service (CRIS #5062-21220-025-000D) and the European Union Framework Programme for Research and Innovation Horizon 2020 (2014–2020) under the Marie Curie Skłodowska Grant Agreement Nr. 847548 for funding. The authors thank all members of the Banfield Laboratory for discussions, in particular Clemence Rodney for careful reading of the manuscript. The authors also thank Abbas Maqbool (JIC Biophysical Analysis), Carlo Martins and Gerhard Saalbach (JIC Proteomics), Julia Mundy (JIC Structural Biology), Phil Robinson (Scientific Photographer), Matt Smoker and Jodie Taylor (TSL Tissue Culture and Transformation), Mark Youles and Liam Egan (TSL SynBio) for their expert help and advice in this work.

## Author Contributions

I.S., H.J.B., M.J.M., J.C.D.C. and M.J.B. designed the experiments; I.S., H.J.B., J.W.B., A.H.C.L., I.H, M.J.M. and J.C.D.C. performed the experiments; I.S., H.J.B., J.W.B., A.H.C.L., I.H., M.J.M., J.C.D.C. and M.J.B. wrote and reviewed the manuscript.

