## Supplementary Information for "Engineering an Exo70 integrated domain of a barley NLR for improved blast resistance"

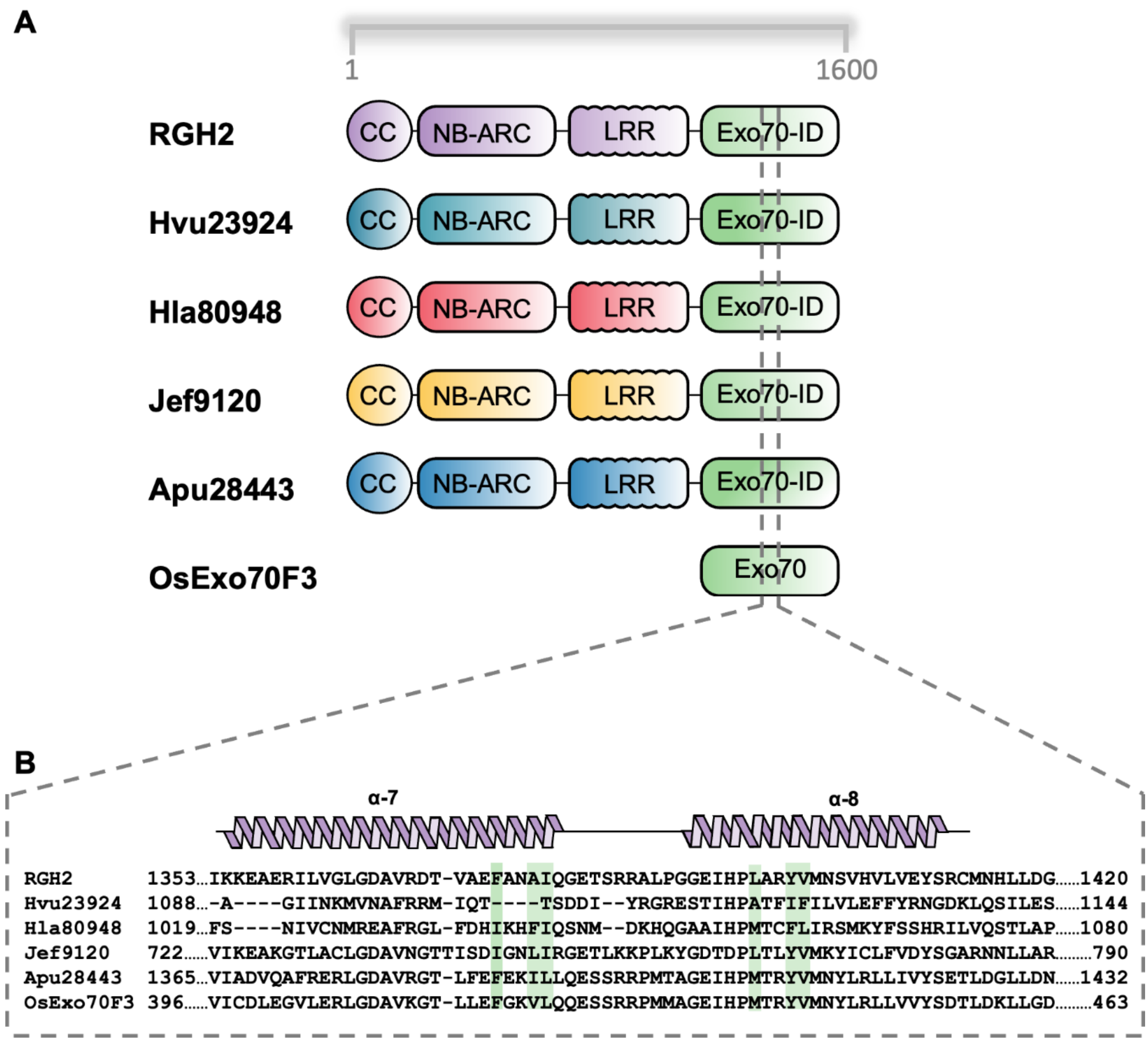


**Supplementary Fig. S1: Sequence alignment of NLRs from different grass species with an Exo70F3 integrated domain.** A. Graphical representation of NLRs with integrated Exo70 domains RGH2, Hvu23924, Hla80948, Jef9120, Apu28443 and OsExo70F3 protein. All NLRs have a CC, NB-ARC and LRR domain. B. Sequence alignment of residues located within OsExo70F3 α-helices 7 and 8 which contribute to the binding interface with AVR-Pii[29] with the equivalent positions of the NLR Exo70 integrated domains. The secondary structure elements are shown above the sequence. Residues that are important for the formation of the AVR-Pii binding pocket are highlighted in green.

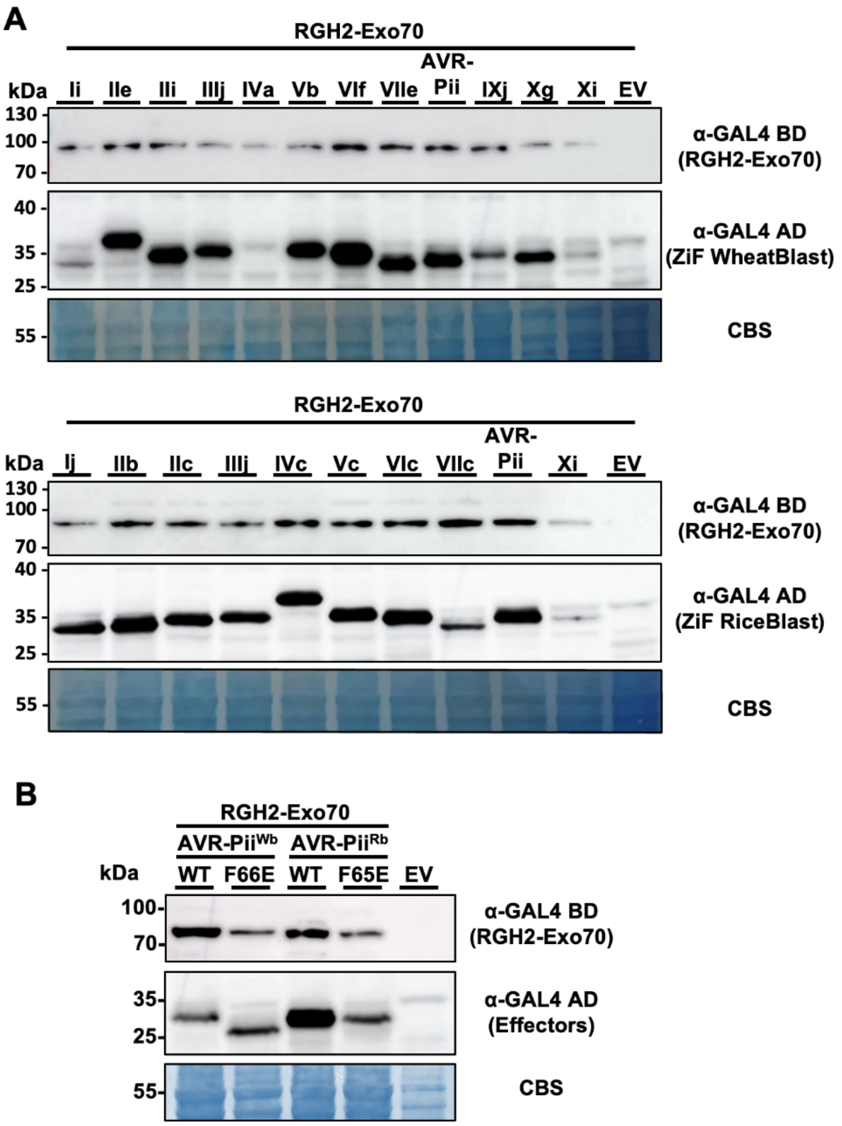


**Supplementary Fig. S2: Protein accumulation in Yeast-Two-Hybrid assays with RGH2-Exo70 domain and Zif effectors shown by Western blot.** A. Anti-GAL4 binding domain (BD) antibodies were used to investigate the presence of RGH2-Exo70 domain in yeast lysate, while anti-GAL4 DNA activation domain (AD) antibodies were used to assess the accumulation of selected rice and wheat blast ZiF effectors. RGH2-Exo70 constructs were only detected weakly, and not all effectors were detected uniformly. Total protein extracts were stained with Coomassie Blue Stain (CBS). B. Same as in A. Anti-GAL4 binding domain (BD) antibodies were used to investigate the presence of RGH2-Exo70 in yeast lysate, while anti-GAL4 DNA activation domain (AD) antibodies were used to assess the accumulation of AVR-Pii$^{Rb}$, AVR-Pii$^{Wb}$, and their mutants. Total protein extracts were stained with Coomassie Blue Stain (CBS).

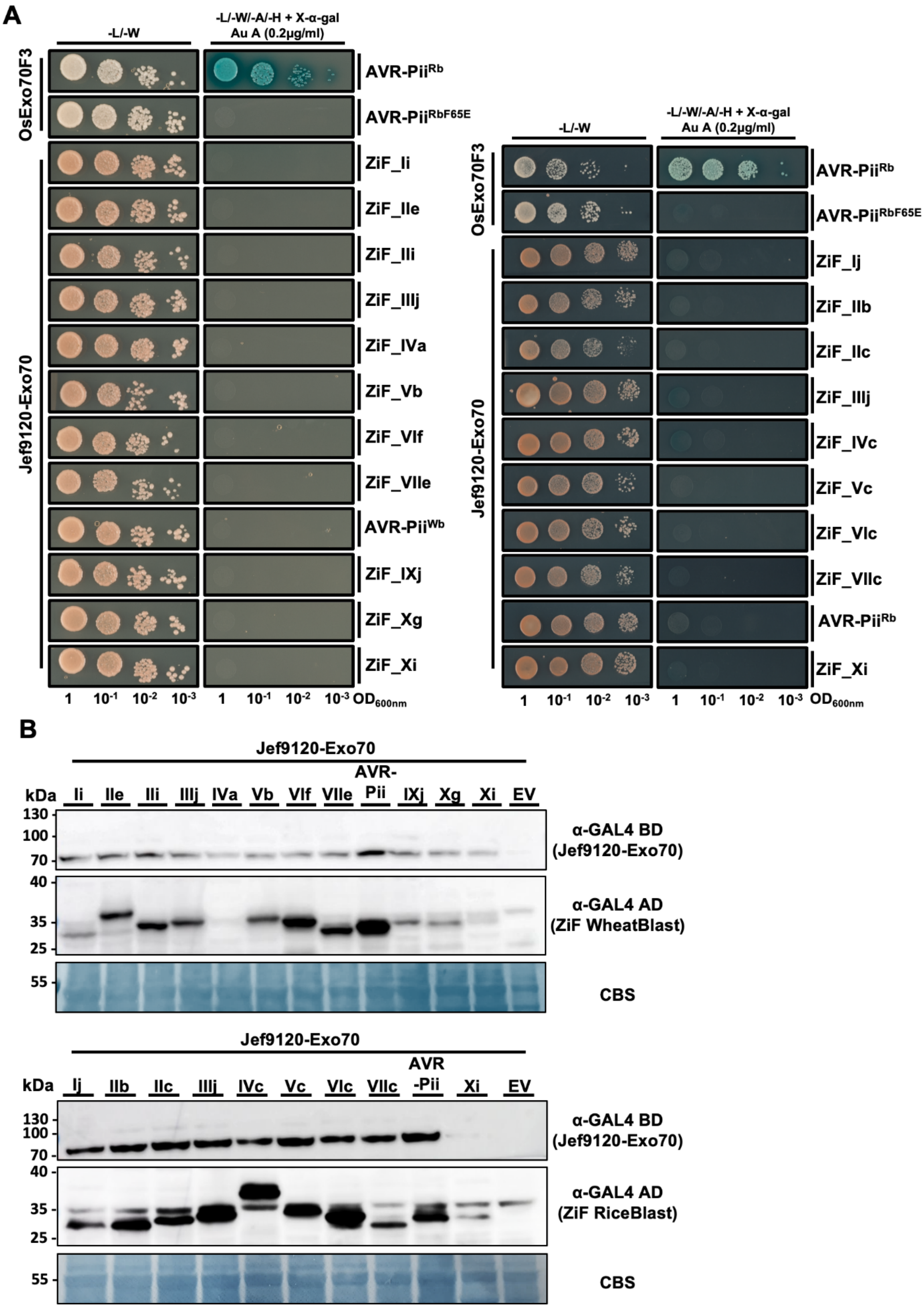


**Supplementary Fig. S3: Protein accumulation in Yeast-Two-Hybrid assays with the Jef9120-Exo70 domain and Zif effectors shown by Western blot.** A. Y2H binding assay of selected ZiF effectors to the Jef9120-Exo70 domain. ZiF effectors were fused to the GAL4 activator domain and co-expressed in yeast cells with

Jef9120-Exo70 domain fused to the GAL4 DNA binding domain. AVR-Pii$^{Rb}$ and AVR-Pii$^{RbF65E}$ binding to OsExo70F3 were used as positive and negative controls, respectively. Yeast cells containing both plasmids were spotted onto selective synthetic dropout (SD) media. For plates with quadruple-dropout media, X-α-gal and aureobasidin A (Au A) were added. Yeast growth was monitored 4 days later. Growth and development of blue coloration indicates protein-protein interactions. The experiment was repeated three times with comparable results. B. Anti-GAL4 binding domain (BD) antibodies were used to determine the presence of the Jef9120-Exo70 domain in yeast lysate, while anti-GAL4 DNA activation domain (AD) antibodies were used to assess the accumulation of ZiF effectors. Exo70 constructs were only detected weakly, and not all effectors were detected uniformly. Total protein extracts were stained with Coomassie Blue Stain (CBS).

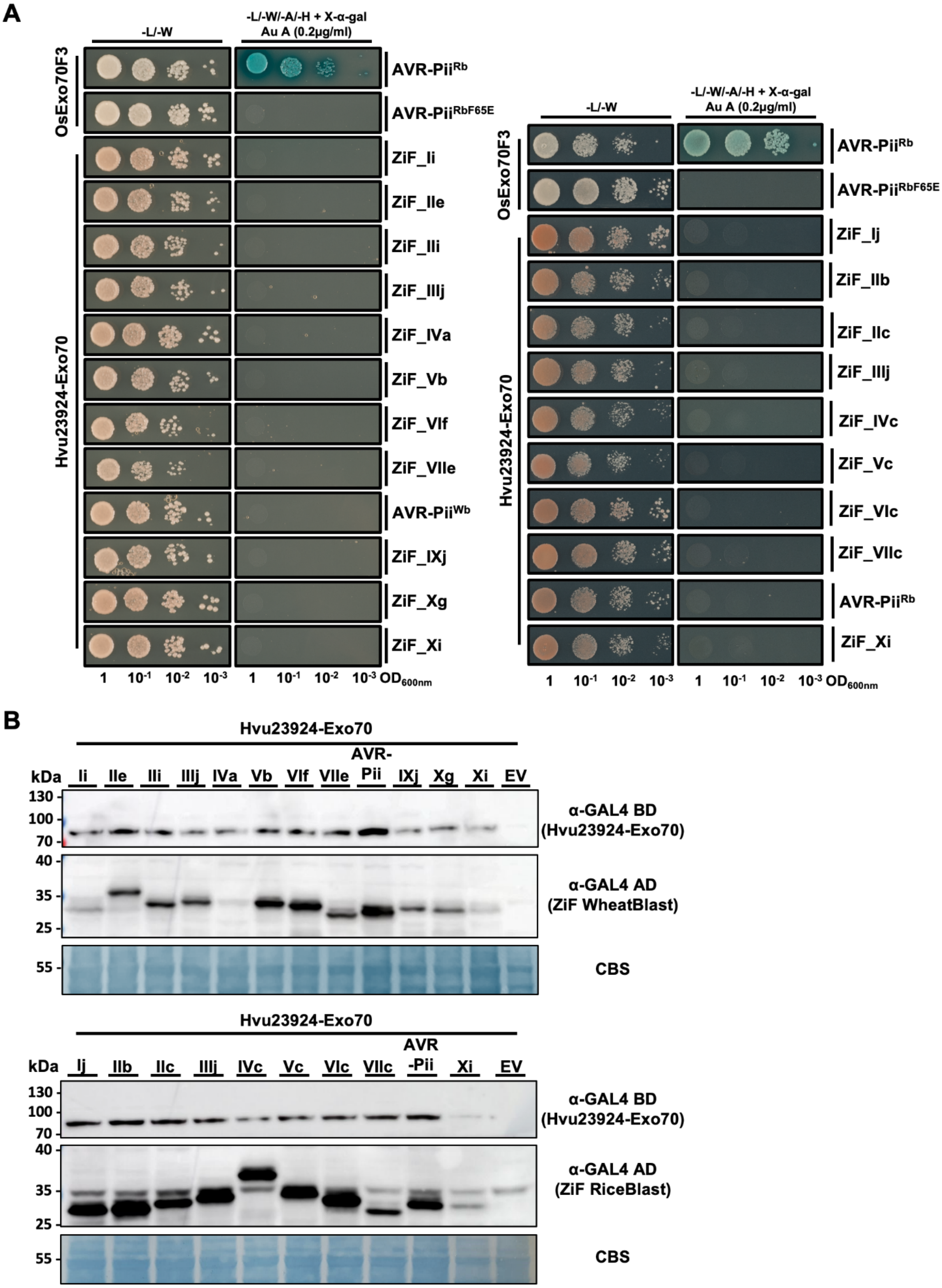


**Supplementary Fig. S4: Protein accumulation in Yeast-Two-Hybrid assays with the Hvu23824-Exo70 domain and Zif effectors shown by Western blot.** A. Y2H binding assay of selected ZiF effectors to the Hvu23824-Exo70 domain. ZiF effectors were fused to the GAL4 activator domain and co-expressed in yeast cells with

Hvu23824-Exo70 domain fused to the GAL4 DNA binding domain. AVR-Pii$^{Rb}$ and AVR-Pii$^{RbF65E}$ binding to OsExo70F3 were used as positive and negative controls, respectively. Yeast cells containing both plasmids were spotted onto selective synthetic dropout (SD) media. For plates with quadruple-dropout media, X-α-gal and aureobasidin A (Au A) were added. Yeast growth was monitored 4 days later. Growth and development of blue coloration indicates protein-protein interactions. The experiment was repeated three times with comparable results. B. Anti-GAL4 binding domain (BD) antibodies were used to determine the presence of the Hvu23824-Exo70 domain in yeast lysate, while anti-GAL4 DNA activation domain (AD) antibodies were used to assess the accumulation of ZiF effectors. Exo70 constructs were only detected weakly, and not all effectors were detected uniformly. Total protein extracts were stained with Coomassie Blue Stain (CBS).

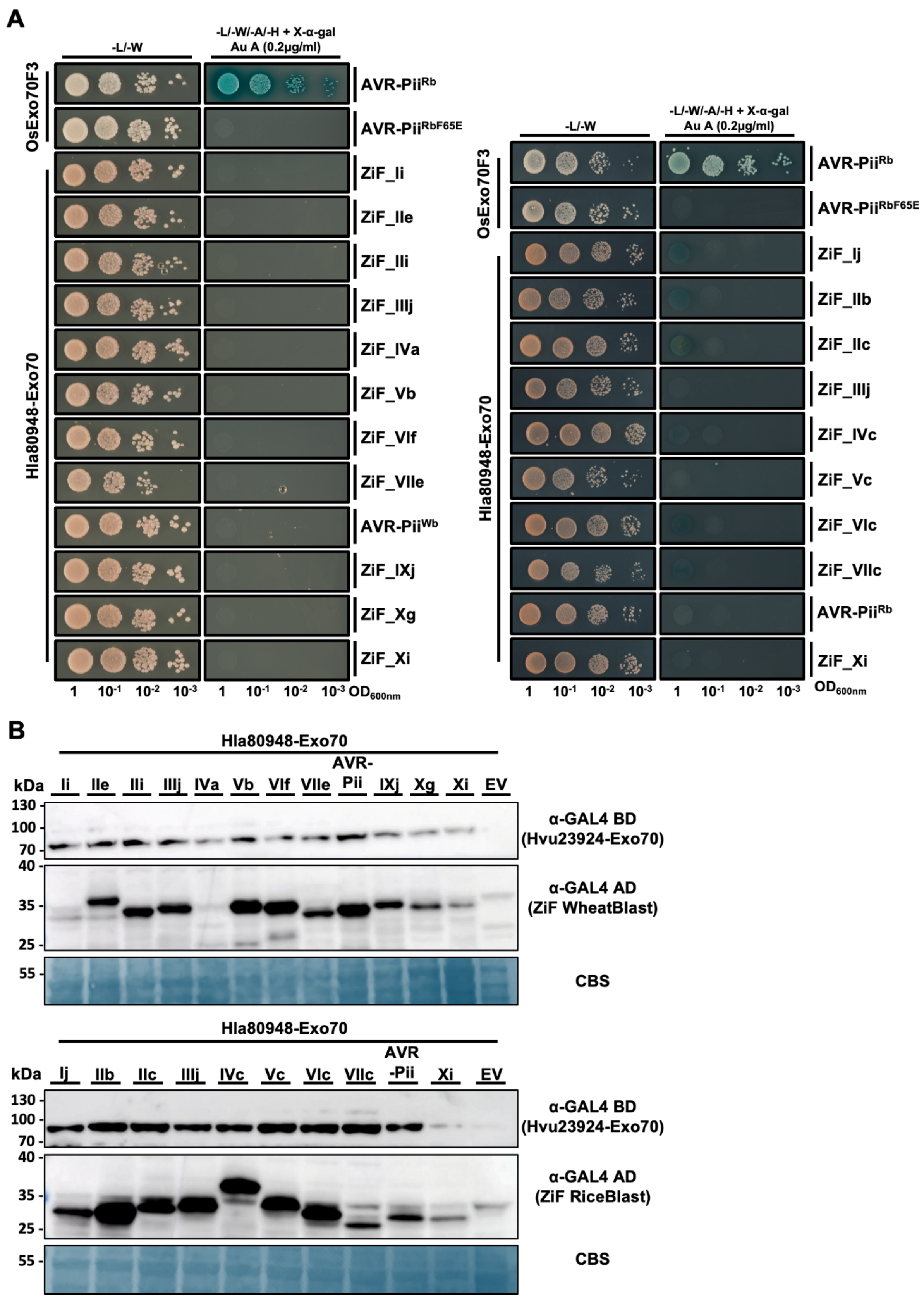


**Supplementary Fig. S5: Protein accumulation in Yeast-Two-Hybrid assays with the Hla80948-Exo70 domain and Zif effectors shown by Western blot.** A. Y2H binding assay of selected ZiF effectors to the Hla80948-Exo70 domain. ZiF effectors were fused to the GAL4 activator domain and co-expressed in yeast cells with

Hla80948-Exo70 domain fused to the GAL4 DNA binding domain. AVR-Pii$^{Rb}$ and AVR-Pii$^{RbF65E}$ binding to OsExo70F3 were used as positive and negative controls, respectively. Yeast cells containing both plasmids were spotted onto selective synthetic dropout (SD) media. For plates with quadruple-dropout media, X-α-gal and aureobasidin A (Au A) were added. Yeast growth was monitored 4 days later. Growth and development of blue coloration indicates protein-protein interactions. The experiment was repeated three times with comparable results. B. Anti-GAL4 binding domain (BD) antibodies were used to determine the presence of the Hla80948-Exo70 domain in yeast lysate, while anti-GAL4 DNA activation domain (AD) antibodies were used to assess the accumulation of ZiF effectors. Exo70 constructs were only detected weakly, and not all effectors were detected uniformly. Total protein extracts were stained with Coomassie Blue Stain (CBS).

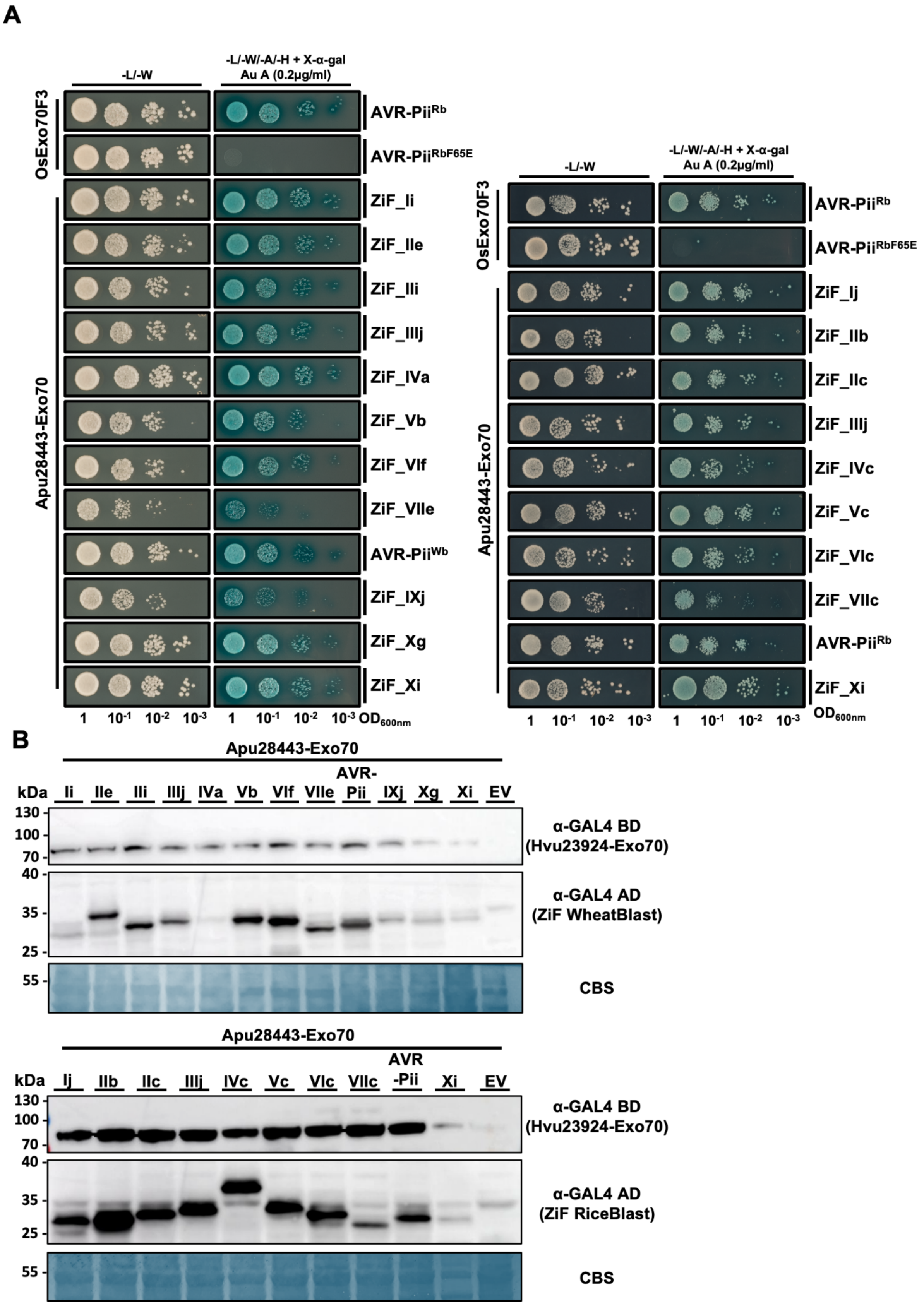


**Supplementary Fig. S6: Protein accumulation in Yeast-Two-Hybrid assays with the Apu28443-Exo70 domain and Zif effectors shown by Western blot.** A. Y2H binding assay of selected ZiF effectors to the Apu28443-Exo70 domain. ZiF effectors were fused to the GAL4 activator domain and co-expressed in yeast cells with Apu28443-Exo70 domain fused to the GAL4 DNA binding domain. AVR-Pii$^{Rb}$ and AVR-Pii$^{RbF65E}$ binding to OsExo70F3 were used as positive and negative controls, respectively. Yeast cells containing both plasmids were spotted onto selective synthetic dropout (SD) media. For plates with quadruple-dropout media, X-α-gal and aureobasidin A (Au A) were added. Yeast growth was monitored 4 days later. Growth and development of blue coloration indicates protein-protein interactions. The experiment was repeated three times with comparable results. B. Anti-GAL4 binding domain (BD) antibodies were used to determine the presence of the Apu28443-Exo70 domain in yeast lysate, while anti-GAL4 DNA activation domain (AD) antibodies were used to assess the accumulation of ZiF effectors. Exo70 constructs were only detected weakly, and not all effectors were detected uniformly. Total protein extracts were stained with Coomassie Blue Stain (CBS).

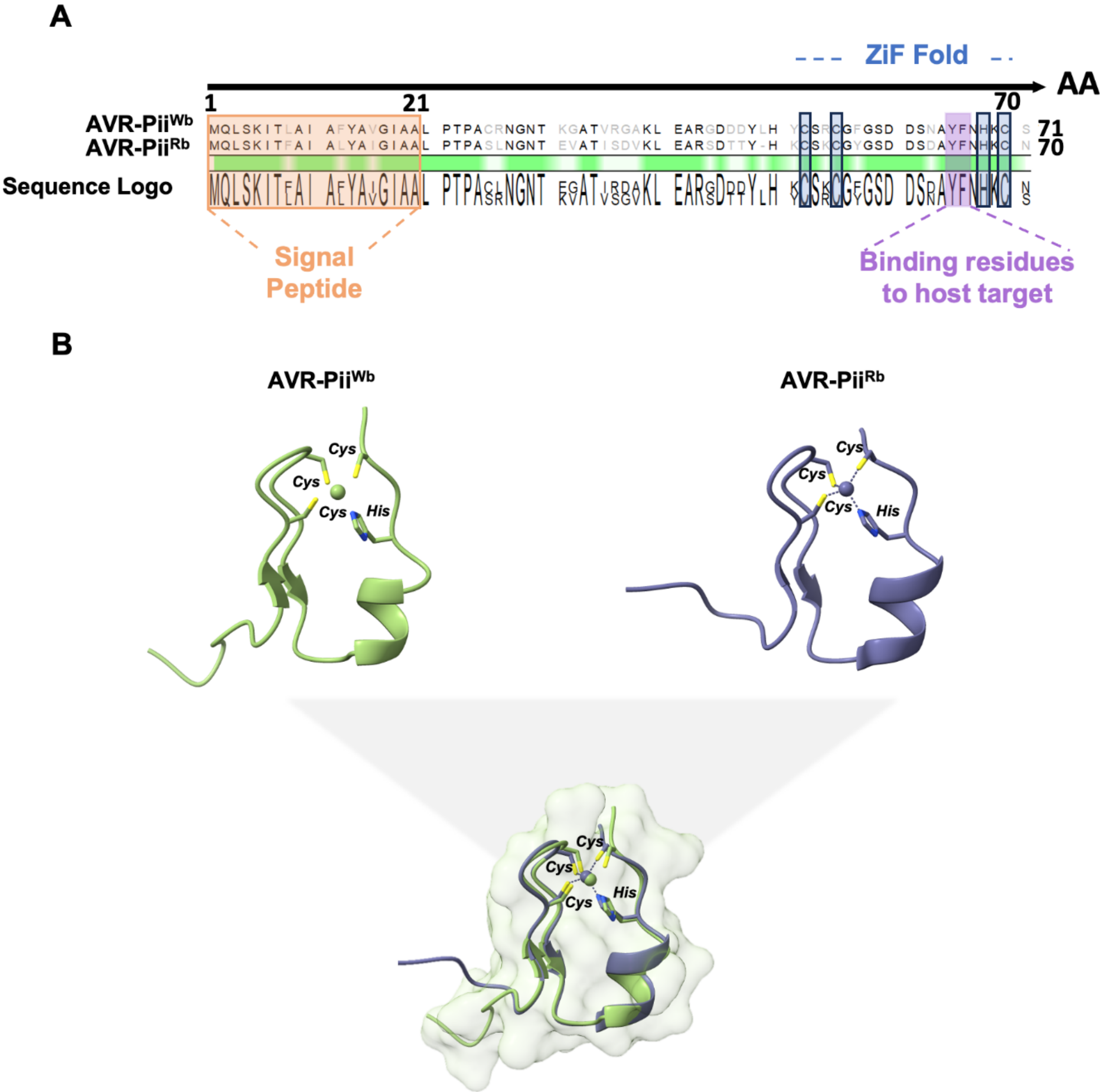


**Supplementary Fig. S7:** A. Amino acid sequence alignment of wheat and rice blast AVR-Pii. Conserved residues are indicated green below the alignment and with the sequence Logo. Orange marks the signal peptide (SP). Purple marked are the residues crucial for host binding. Blue indicates residues CCHC that represent the Zinc finger fold (ZiF). B. Structural comparison between rice AVR-Pii$^{Rb}$ (PDB: 7PP2) and AVR-Pii$^{Wb}$ predicted by Alphafold3[50] and superimposition of the two effectors, respectively.

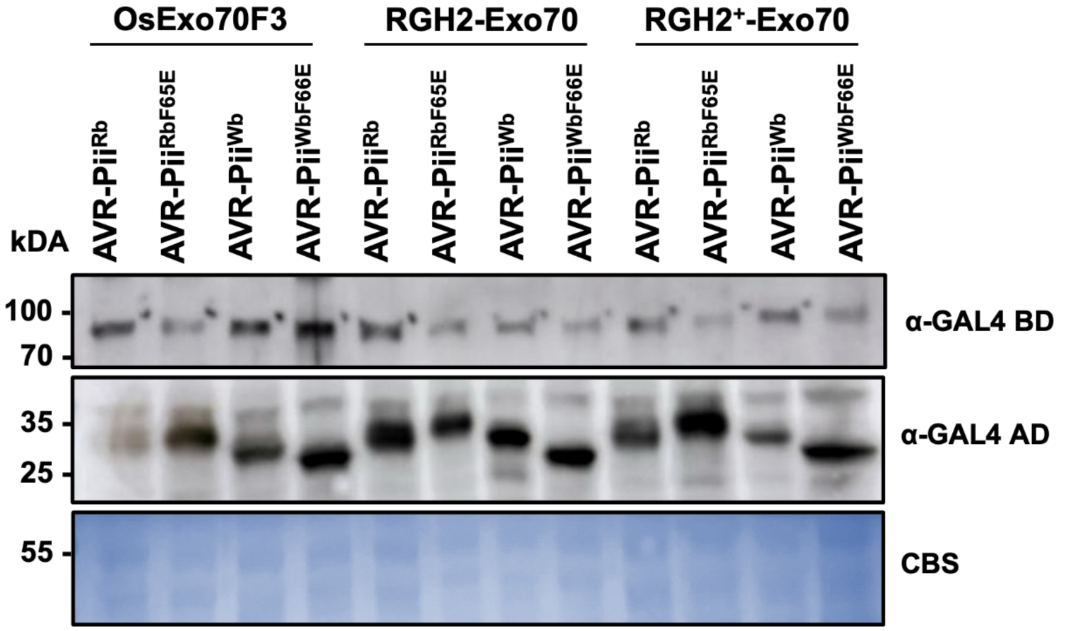


**Supplementary Fig. S8: Protein accumulation in Yeast-Two-Hybrid assays with OsExo70F3, RGH2-Exo70 domain, $RGH2^{+}$-Exo70 domain and blast AVR-Pii variants analysed by Western blot.** Anti-GAL4 binding domain (BD) antibodies were used to determine the presence of OsExo70F3, and the RGH2-Exo70 and $RGH2^{+}$-Exo70 domains in yeast lysate, while anti-GAL4 DNA activation domain (AD) antibodies were used to assess the accumulation of effectors and respective mutants. Total protein extracts were stained with Coomassie Blue Stain (CBS).

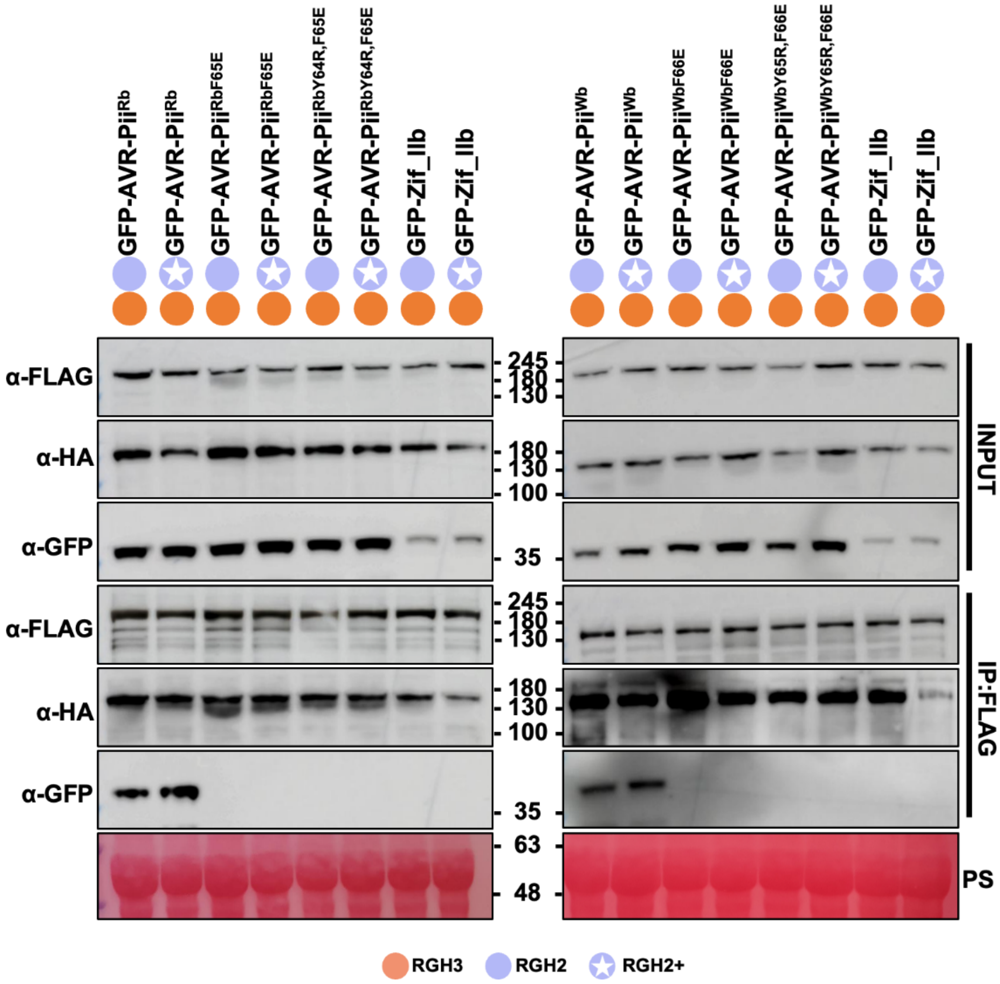


**Supplementary Fig. S9: Extended data for Co-IP experiments shown in Fig. 3D.** Agroinfiltration of *N. benthamiana* leaves was used to co-express sensor NLR FLAG-RGH2 or FLAG-RGH2$^{+}$ and the helper NLR RGH3-HA with GFP-AVR-Pii variants and mutants or ZiF_IIb (negative control) under the *MASpro* promoter. Pulldowns were performed using an anti-FLAG antibody. Immunoblot analysis with an anti-GFP antibody showed that FLAG-RGH2 and FLAG-RGH2$^{+}$ copurified with the GFP-AVR-Pii variants but not with the mutants or ZiF_IIb. RGH2 and RGH2$^{+}$ were only detected weakly in the inputs, but immunoprecipitated proteins are clear. Total protein extracts were stained with Ponceau (PS). Sections of the blots are shown in Fig. 3. The experiment was repeated three times with similar results.

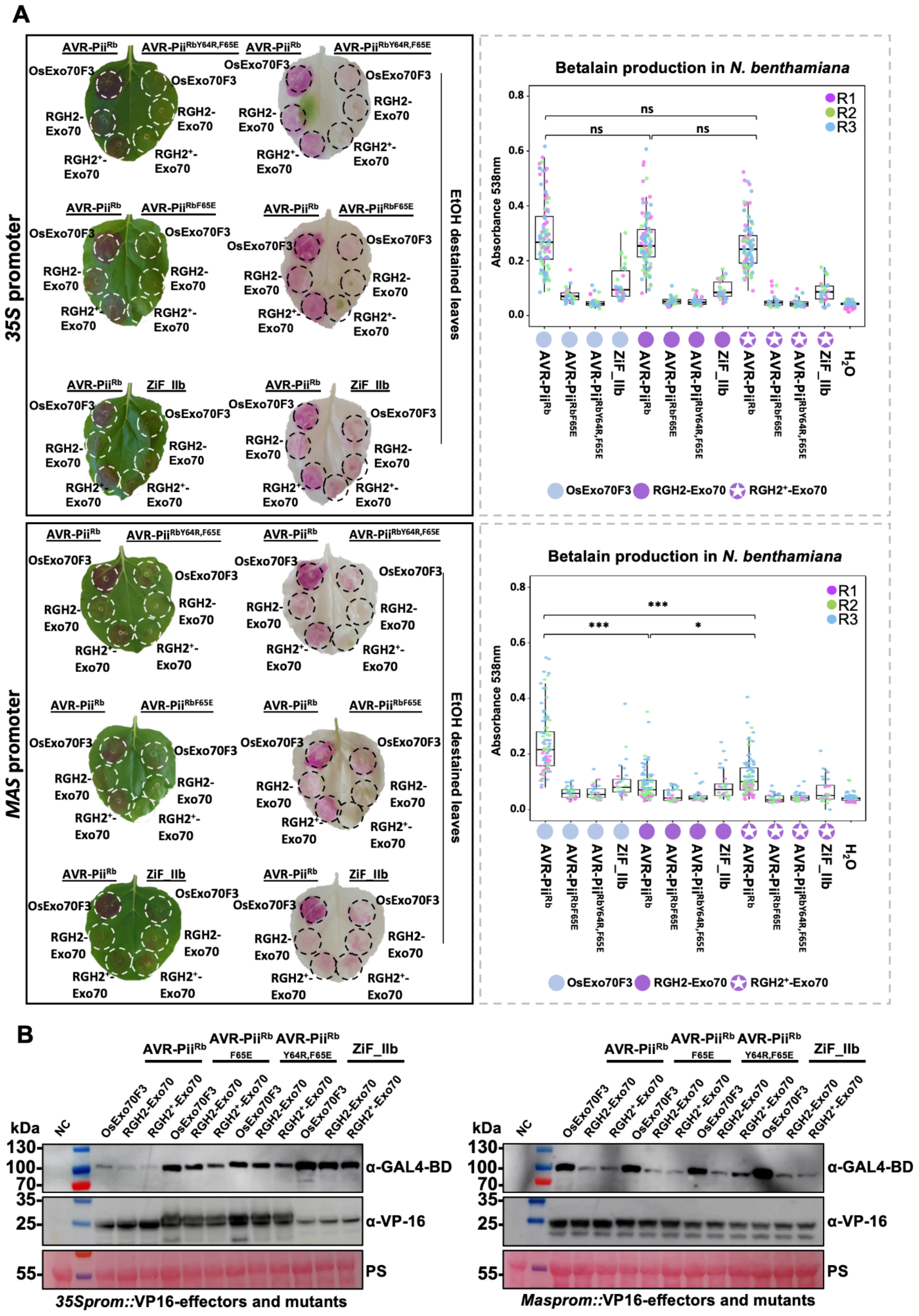


**Supplementary Fig. S10: Extended data for split GAL4 Ruby assay shown in Fig. 4A and 4B for AVR-Pii$^{Rb}$.** A. Agroinfiltration of *N. benthamiana* leaves was used to co-express OsExo70F3, RGH2-Exo70 or RGH2$^{+}$-Exo70 with VP16-AVR-Pii$^{Rb}$, VP16-AVR-Pii$^{RbF65E}$, VP16-AVR-Pii$^{RbY64R,F65E}$ or effector VP16-ZiF_IIb, under the control of the *35Spro* or *MASpro* promoter as indicated. Betalain production was assessed 3 days post infiltration. Box plots represent the median (horizontal line), upper and lower quartiles (boxes), and 1.5× interquartile range (whiskers). Asterisks indicate statistically significant differences from the control (NS: not significant, *$p$< 0.01, ***$p$< 0.001, one-way ANOVA followed by Tukey's test). Replicates are indicated with different colours (magenta-Replicate 1, green-Replicate 2, blue-Replicate. The experiment was repeated three times with similar results. 3). B. Accumulation of OsExo70F3, RGH2-Exo70 and RGH2$^{+}$-Exo70 under the control of the *35Spro* promoter in leave tissue was assessed by western blot with anti-GAL4-BD antibody. Accumulation of effectors under the control of either the *35Spro* (left) or the *MASpro* (right) promoter were assessed by western blot with anti-VP16 antibody. Weak accumulation of some Exo70 proteins was observed in western blots (left panel). Total protein extracts were stained with Ponceau (PS).

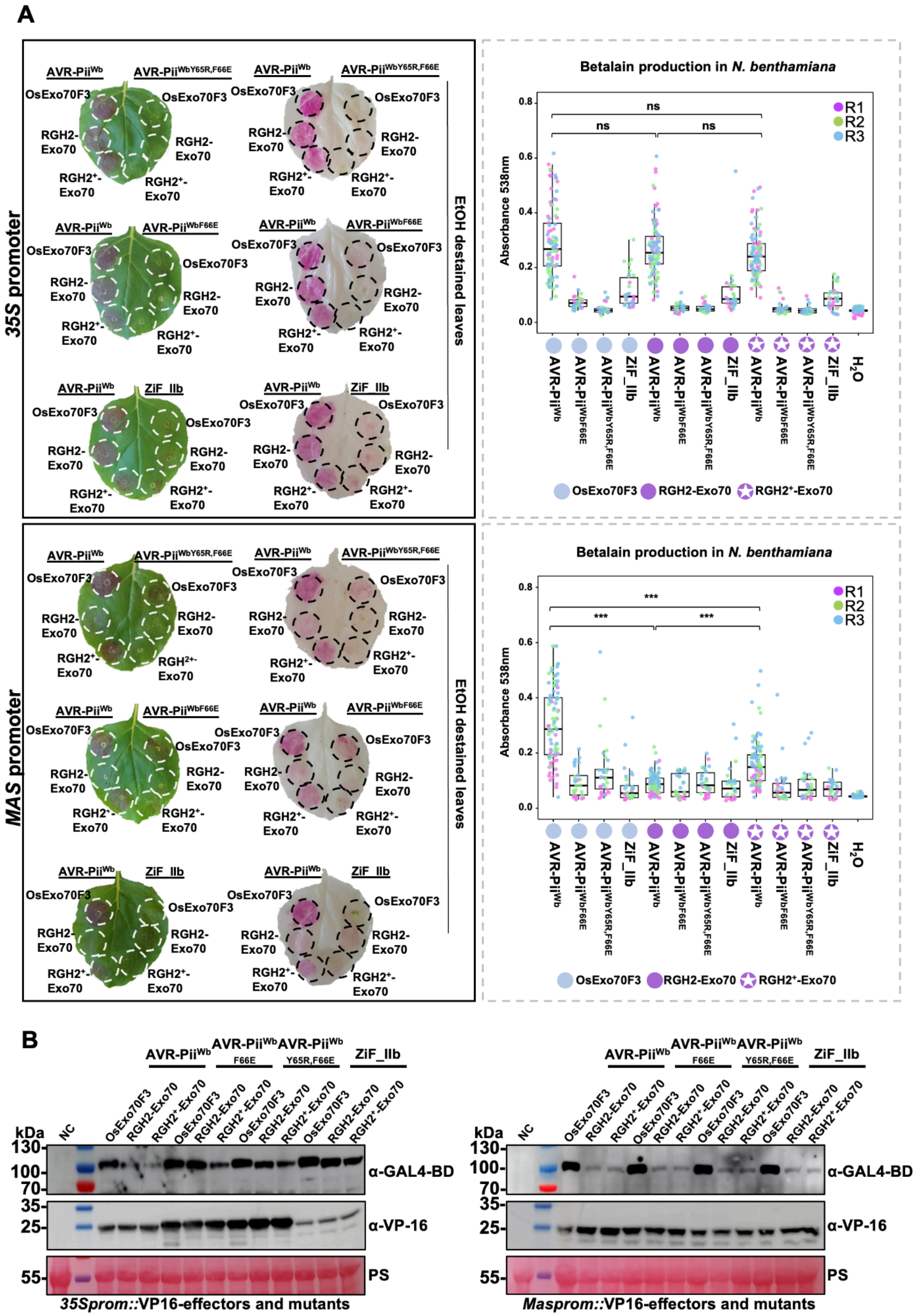


**Supplementary Fig. S11: Extended data for split GAL4 Ruby assay shown in Fig. 4A and 4B for AVR-Pii$^{Wb}$.** A. Agroinfiltration of *N. benthamiana* leaves was used to co-express OsExo70F3, RGH2-Exo70 or RGH2$^{+}$-Exo70 with VP16-AVR-Pii$^{Wb}$, VP16-AVR-Pii$^{WbF65E}$, VP16-AVR-Pii$^{WbY64R,F65E}$ or effector VP16-ZiF_IIb, under the control of the *35Spro* or *MASpro* promoter as indicated. Betalain production was assessed 3 days post infiltration. Box plots represent the median (horizontal line), upper and lower quartiles (boxes), and 1.5× interquartile range (whiskers). Asterisks indicate statistically significant differences from the control (NS: not significant, ***$p < 0.001$, one-way ANOVA followed by Tukey's test). Replicates are indicated with different colours (magenta-Replicate 1, green-Replicate 2, blue-Replicate 3). The experiment was repeated three times with similar results. B. Accumulation of OsExo70F3, RGH2-Exo70 and RGH2$^{+}$-Exo70 under the control of the *35Spro* promoter in leave tissue was assessed by western blot with anti-GAL4-BD antibody. Accumulation of effectors under the control of either the *35Spro* (left) or the *MASpro* (right) promoter were assessed by western blot with anti-VP16 antibody. Weak accumulation of some Exo70 proteins was observed in western blots (left panel). Total protein extracts were stained with Ponceau (PS).

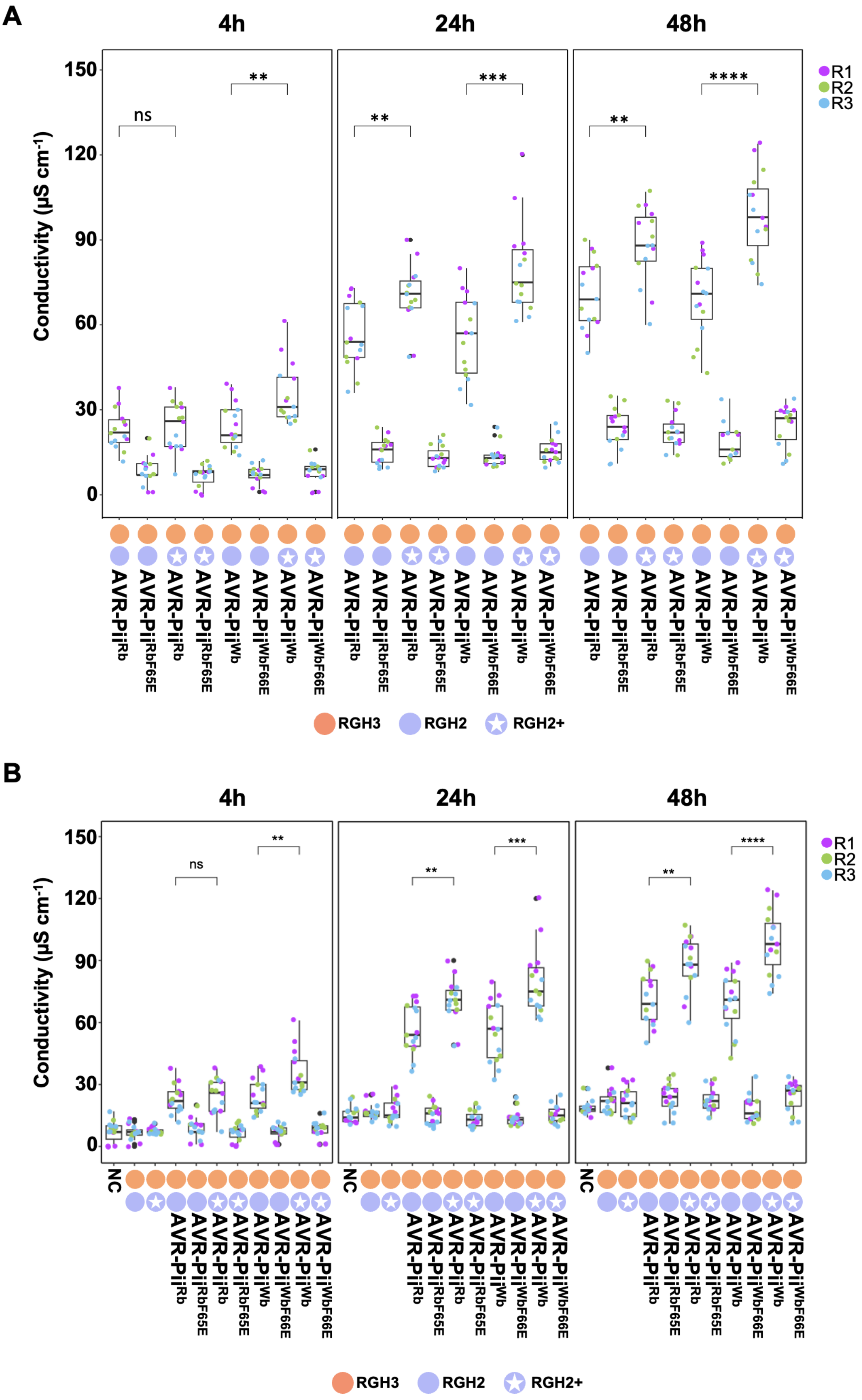


**Supplementary Fig. S12: Extended Split Ion leakage assay for Fig. 5B.** A. Conductivity (µS cm-1) of *N. benthamiana* leaf tissue infiltrated with RGH2/RGH3 or RGH2$^{+}$/RGH3 NLR pairs with GFP-AVR-Pii variant or mutants at 4, 24, and 48 hours post infiltration (hpi). Box plots represent the median (horizontal line), upper and lower quartiles (boxes), and 1.5× interquartile range (whiskers). Statistically significant differences are denoted by an asterisk (NS: not significant, **$p$<0.01, ***$p$<0.001, ****$p$<0.0001 Student's t-test). Data from three independent experimental replicates are presented (n=5 plants per experiment). B. Same as Supplementary Fig. S12A with additional controls. Additional controls include RGH2/RGH3 and RGH2+/RGH3 co-infiltrated together and only buffer infiltration (NC – negative control). Replicates are designated with different colours (magenta-Replicate 1, green- Replicate 2, blue-Replicate 3).
